# Different factors drive community assembly of rare and common ectomycorrhizal fungi

**DOI:** 10.1101/2022.04.06.487384

**Authors:** Laura G. van Galen, David A. Orlovich, Janice M. Lord, Julia Bohorquez, Andy R. Nilsen, Tina C. Summerfield, Matthew J. Larcombe

## Abstract

Understanding what drives community assembly processes and how communities respond to environmental gradients are fundamental goals in community ecology. Ectomycorrhizal fungi support major forest systems across the globe, but the diversity, distribution and environmental controls affecting ectomycorrhizal community composition are unknown in many regions, particularly in the southern hemisphere. Here we investigate the assembly of ectomycorrhizal fungal communities based on eDNA samples from 81 *Nothofagus* forests across New Zealand’s South Island. We apply zeta diversity analysis and multi-site generalised dissimilarity modelling (MS-GDM) to investigate assembly patterns and quantify the effects of 43 biotic and environmental variables on community turnover. The zeta diversity MS-GDM framework differentiates between the environmental factors driving turnover of rare and common species, so provides a more complete picture of community dynamics than traditional beta diversity analyses. Results showed that community assembly was dominated by deterministic rather than stochastic processes. Soil variables were important drivers across the full range of rare, intermediate and common species. Ground cover variables, forest patch size and rainfall had greater effects on turnover of rare species, whereas temperature variables and host tree size had greater effects on common species turnover. Applying these methods for the first time to fungi demonstrates that there are distinct differences in the ecological processes affecting different aspects of the ectomycorrhizal community, which has important implications for understanding the functional effects of community responses to environmental change.

## 2. Introduction

The structure of ecological communities is governed by a combination of deterministic niche-based processes and stochasticity (Hanson *et al*. 2012; Stegen *et al*. 2013; Vellend 2010). Whilst biotic and abiotic factors can have strong effects on species presence and abundance through deterministic selection, communities are also influenced to varying degrees by stochastic processes such as dispersal, ecological drift and speciation (Vellend 2010). Understanding the degree to which these factors influence the structure and function of communities is a central goal in ecology and biogeography, and is fundamental for understanding how communities respond to environmental change (Bahram *et al*. 2015; Hanson *et al*. 2012).

Understanding microbial community assembly processes is particularly important because microbial communities underpin the structure and function of all ecosystems on Earth (Hanson *et al*. 2012; Peay and Bruns 2014). Ectomycorrhizal symbiotrophs have particularly strong effects on terrestrial ecosystems, as they facilitate the exchange of nutrients and other resources that support major temperate and tropical forest systems across the globe (Martin *et al*. 2016). Ectomycorrhizal fungi are taxonomically and evolutionarily diverse (Rinaldi *et al*. 2008), forming at least 78 independently evolved lineages within the phyla Basidiomycota, Ascomycota and Mucoromycota (Tedersoo and Smith 2013). Taxa also display a wide range of life history strategies and morphological and functional types that likely respond to environmental conditions in different ways (Koide *et al*. 2014; Martin *et al*. 2016; van der Heijden *et al*. 2015). Whilst research into soil microbiomes has expanded since the development of eDNA techniques, the diversity and structure of ectomycorrhizal communities, and the factors driving species turnover, still remain largely unknown in many regions where large-scale data are not yet available (Peay *et al*. 2010).

A number of well-developed approaches exist for investigating the relative importance of selection, dispersal, drift and speciation acting on particular communities, and for exploring effects of different environmental variables. For example, the null model approach by Stegen *et al*. (2013) and Stegen *et al*. (2015) uses phylogenetic turnover between communities to understand ecological selection processes, and techniques such as generalised dissimilarity modelling (GDM; Ferrier *et al*. 2007) can be used to determine the influence of environmental variables on spatial community turnover patterns. However, these techniques focus on turnover of the species assemblage between pairs of communities or sites (i.e., beta diversity), which does not capture all components of assemblage partitioning, such as diversity shared between three or more sites (Hui and McGeoch 2014; Latombe *et al*. 2017; Latombe *et al*. 2019). Because rare species are less likely to be shared between two sites than common species, differences between site pairs (beta diversity) disproportionally represents turnover in rare species, and underestimates the contribution of more widespread species to turnover (Latombe *et al*. 2017; Latombe *et al*. 2018a). Characterising turnover patterns using “zeta diversity”, the average number of shared species between groups of *n* (two or more) sites, has been proposed as a way to overcome these issues (Hui and McGeoch 2014). The value of *n* (i.e., the number of sites compared) is termed the “zeta order”, and assessing how patterns of zeta diversity change with increasing zeta order can provide insight into the processes acting upon both rare species (patterns at lower zeta orders) and common species (patterns at higher zeta orders) (Hui and McGeoch 2014; Latombe *et al*. 2017; McGeoch *et al*. 2019).

Different patterns in the parametric form of decline in zeta diversity with increasing zeta order can show the degree to which stochastic and deterministic processes drive turnover (McGeoch *et al*. 2019; see Section 3.6.1 for details). When deterministic processes are important, multisite generalised dissimilarity modelling (MS-GDM) can be used to examine environmental drivers of species turnover (Latombe *et al*. 2017). MS-GDM is an extension of traditional GDM that uses zeta diversity to assess turnover patterns, allowing relative effects of environmental variables influencing both rare and more common species turnover to be investigated (Latombe *et al*. 2017; Latombe *et al*. 2018b). To date, zeta diversity patterns and MS-GDM have been used to investigate patterns of community assembly and environmental gradients in plants (e.g., Hui *et al*. 2018; Latombe *et al*. 2018b; Lazarina *et al*. 2019), invertebrates (e.g., Baird *et al*. 2019; Cai *et al*. 2021; Latombe *et al*. 2019; Simons *et al*. 2019), parasites (e.g., Krasnov *et al*. 2020a; Krasnov *et al*. 2020b), birds (Ascensão *et al*. 2020; Latombe *et al*. 2017), fish (Pettersen *et al*. 2021), amphibians (Fonte *et al*. 2021), and bacteria (Bay *et al*. 2020).

Previous ectomycorrhizal research shows that these fungal communities are affected by a combination of stochastic and deterministic assembly processes, and the relative importance of each depends on the scale and region examined (Bahram *et al*. 2013; Beck *et al*. 2015; Chen *et al*. 2018; Glassman *et al*. 2017; Kranabetter *et al*. 2015; Taylor *et al*. 2014). Numerous environmental variables have been identified as important drivers of species turnover, including soil pH, nitrogen availability, organic matter, elevation, temperature and precipitation (Bahram *et al*. 2012; Glassman *et al*. 2017; Kranabetter *et al*. 2015; Nouhra *et al*. 2013; Suz *et al*. 2014; Taylor *et al*. 2014; Tedersoo *et al*. 2012a; van der Linde *et al*. 2018). Many ectomycorrhizal taxa also show strong preference for certain host lineages, and variation in host traits can affect community composition (Tedersoo *et al*. 2008; van der Linde *et al*. 2018). Past research has focussed on northern hemisphere systems (although see Marion *et al*. 2021; Nouhra *et al*. 2013; Tedersoo *et al*. 2008), and has been predominantly restricted to beta diversity analyses of pairwise turnover and so heavily influenced by rare species’ patterns. To the best of our knowledge, zeta diversity analysis has not previously been applied to fungal communities, and so whether patterns influencing more widespread species differ to those influencing rare species is unknown. Current understanding of the functional roles of different ectomycorrhizal species is predominantly restricted to more common taxa that are easily detectable (Tedersoo and Smith 2013). Therefore, quantifying factors affecting more widespread species will improve our capacity to understand the functional implications of how communities respond to environmental variation (Koide *et al*. 2014).

This study applies the zeta diversity MS-GDM framework to ectomycorrhizal fungal communities in New Zealand, to investigate ectomycorrhizal fungal community assembly processes and assess how assembly patterns and environmental drivers of rare and more widespread species differ. We use eDNA techniques to characterise ectomycorrhizal fungal communities at 81 *Nothofagus* (southern beech, Nothofagaceae) forest sites spanning the length and breadth of New Zealand’s c. 150,000 km^2^ South Island and test the response of communities to 43 biotic and environmental variables collected at the same locations. This region is dominated by *Nothofagus* forest spanning a relatively wide range of elevations, climatic conditions, forest structures, forest patch sizes, and soil types, and includes stands of all five of New Zealand’s *Nothofagus* species (*N. menziesii, N. fusca, N. truncata, N. solandri* var. *solandri*, and *N. solandri* var. *cliffortioide*s) (Ogden *et al*. 1996). This allows the relative effects of host species versus other environmental variables on ectomycorrhizal turnover to be examined. Large-scale below-ground ectomycorrhizal fungal data has not previously been available for New Zealand or other southern hemisphere regions.

Specifically, we aim to 1) investigate the extent to which stochastic and deterministic processes drive ectomycorrhizal community assembly; 2) examine the relative effects of host species, soil properties, forest structure, ground covers, topography and climatic variables on fungal community turnover; and 3) explore differences in factors driving community turnover of rare versus common species.

## 3. Methods

### 3.1 Site Selection

Eighty-one survey sites were established in *Nothofagus* forests across southern New Zealand (Figure 1). To select sites, spatially-distributed target locations and nearby backup locations were identified using the EcoSat Forest digital mapping layer of indigenous New Zealand forest classes (Landcare Research 2014), along with local knowledge of smaller forest patches not included in the mapping layer. Target locations were assessed in the field, and survey sites were established if they a) were accessible, b) contained forest canopies dominated by *Nothofagus* species, and c) were not in close proximity to other ectomycorrhizal fungal host species (e.g., *Leptospermum, Kunzea, Eucalyptus*, or ectomycorrhizal conifer species). If these criteria were not met, the nearby backup locations were used instead. Sites were at least 4 km apart.

**Figure 1:**
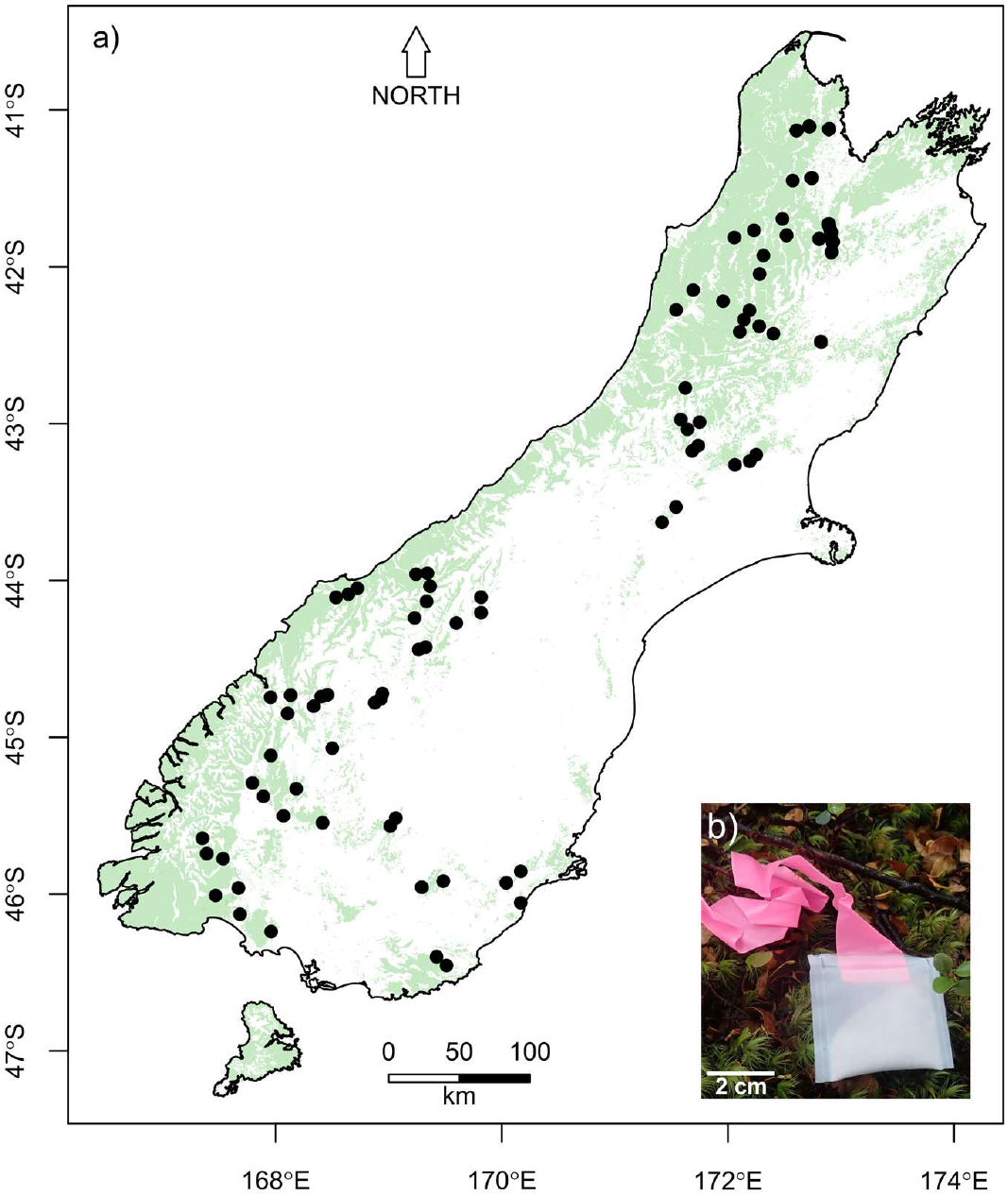
a) locations of the 81 survey sites within the South Island of New Zealand. Green shading is native forest cover (including non-*Nothofagus* forest). The inset b) shows one of the 4,050 hyphal ingrowth bags that were buried across the sites to sample fungal diversity. The gap in sites along the central west coast reflects the “Westland beech gap” that is devoid of *Nothofagus* forest (Heenan *et al*. 2017).

### 3.2 Ectomycorrhizal fungi sampling

Ectomycorrhizal fungi were sampled using hyphal ingrowth bags, which allow fungal hyphae to grow inside whilst excluding plant roots, and so allow the relatively clean retrieval of growing mycelium from the soil (Wallander *et al*. 2001). Ingrowth bags also select for mycorrhizal fungi over saprotrophic fungi that rely on substrate carbon sources (Wallander *et al*. 2001). Although ingrowth bags may favour fast-growing species, this method is more comprehensive than other methods such as root-tip sampling, and is well suited for making comparisons between different sites (Wallander *et al*. 2013). We constructed 4 × 4 cm hyphal ingrowth bags from 50 μm nylon mesh, filled them with 5 g of acid-washed sand (mean grain size 105 μm, SE ± 4.5) and then heat-sealed them with flagging tape sealed into the top to facilitate retrieval (Figure 1b). All bags were heat-treated before deployment for 36 hours at 60 °C to reduce the risk of contamination. The following sampling protocol was designed to maximise the capture of hyphae and fungal taxa that have characteristically patchy distributions (Anderson and Cairney 2007). At each of the 81 survey sites, 10 1.5 × 1.5 m plots were established at least 15 m from the forest edge or any roads, tracks, or other disturbed areas, with plots spaced at least 5 m apart. Five hyphal ingrowth bags were buried in each plot spaced 1 to 1.5 m apart (50 bags per site, 4,050 in total). Ingrowth bags were buried horizontally between the organic and mineral soil layer up to a maximum of 15 cm deep, usually between 2 and 10 cm. Bags remained in the ground for approximately nine months between January and November 2019. Upon retrieval, bags were frozen in the field using dry ice, and stored at −20 °C until DNA extraction.

### 3.3 Fungal DNA extraction, amplification and sequencing

Of the 4,050 ingrowth bags buried across the 81 sites, 97.4% (3,943) were recovered. Of these, 3,639 (92.3%) were still buried in the ground upon collection, and 304 bags were found sitting on the surface. The remaining 157 bags were missing, however all sites were represented by at least 40 bags. Bags from all 10 plots were recovered from all but two sites where all bags were missing from one plot at each. The hyphal ingrowth bags retrieved from each plot were pooled for DNA analysis, resulting in 10 samples for most sites and nine for the two sites with missing plots. Therefore, 808 pooled samples from the 81 sites were sequenced.

DNA was extracted from fungal hyphae collected in the hyphal ingrowth bags and sequenced using high-throughput sequencing. Methods to separate the hyphae from the sand of each ingrowth bag were adapted from Branco *et al*. (2013). Specifically, each ingrowth bag was cut open and the contents scraped into a 50 ml centrifuge tube. We added 3 ml of autoclaved ultrapure water, mixed using a vortex mixer for 3 seconds, and let each tube settle for 15 minutes to allow the sand to sink to the bottom. Observations showed hyphae floating on top of the sand and suspended in the water. Using a cut 1,000 μl pipette tip, floating hyphae and liquid from the top of the sand was transferred into a new 50 ml centrifuge tube. Examining the remaining sand under a microscope showed that this method was successful in extracting almost all visible hyphae from the sample. At this point, the hyphae from each of the five bags in each plot was pooled into a single 50 ml centrifuge tube. The hyphal clumps from each pooled sample were then macerated with sterilised scissors (sterilised with a 0.4% bleach solution) to homogenise the sample, and a 1 ml sub-sample was transferred to a 1.5 ml screw-cap tube for DNA extraction. This sub-sample was centrifuged for 2 minutes at 14,100 × g to pellet the hyphae, after which all remaining liquid was removed. To lyse cells prior to DNA extraction, between five and ten 2.3 mm zirconium beads were added to each tube, along with 400 μl of Qiagen DNeasy Plant Mini-Kit DNA extraction buffer AP1 and 3 μl of RNase A (concentration 100 mg/ml), and tubes were shaken on a bead-beater (AlphaTec) for three bursts of 30 seconds, with 30 seconds of rest in between each burst. Samples were then placed in a heated block at 60 °C for 10 minutes, and the DNA extraction process was continued using the Qiagen DNeasy Plant Mini-Kit, following the manufacturer’s instructions.

We undertook PCR amplification of the internal transcribed spacer 2 (ITS2) region using the forward primer fITS7 (Ihrmark *et al*. 2012) and reverse primer ITS4 (White *et al*. 1990), with appropriate linkers for Illumina MiSeq sequencing (Table 1). Each PCR reaction included 12.5 μl of KAPA HiFi HotStart Ready Mix, 1 μl (10 pmol) each of forward and reverse primers, 1 μl of 50 mg/ml bovine serum albumin, 9.5 μl of autoclaved ultrapure water, and 1 μl of undiluted DNA extract. A thermal gradient PCR showed the optimal annealing temperature for the primer pair and linkers was 57 °C, so PCR amplification was carried out using the following protocol: 95 °C for 3 minutes, followed by 30 cycles of 95 °C for 30 seconds, 57 °C for 30 seconds, and 72 °C for 45 seconds, followed by a final extension period of 5 minutes at 72 °C. Negative controls were included for all PCR runs to ensure no contamination occurred. We used gel electrophoresis with a 1.5% agarose in tris-acetic acid-EDTA (TAE) gel stained with ethidium bromide to ensure that amplification of each sample was successful. Between 15 and 20 μl of PCR product for each sample was purified with 1.8 times the sample volume of Agencourt AMPure XP beads, following the manufacturer’s instructions. Concentrations were quantified using a Qubit dsDNA High Sensitivity Assay Kit, after which samples were diluted to equal concentrations. Samples were then sent to Massey Genome Service Sequencing Facility at Massey University, New Zealand, for second-round PCR, library preparation, and Illumina MiSeq sequencing on 2 × 250 bp paired-end runs. Due to the high volume of samples, samples were sequenced across three separate runs, with samples randomly allocated to a run to minimize bias. For two of the runs, a 350 bp size-selection clean-up for the pooled library was carried out to improve library quality.

**Table 1:**
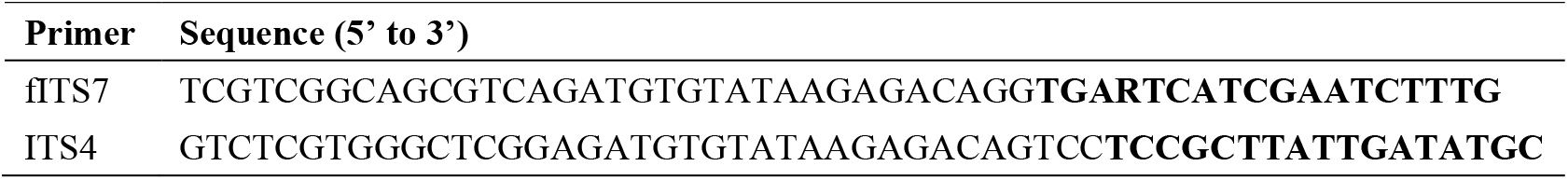
The primers used to sequence the ITS2 region (bold) with associated linkers (non-bold).

### 3.4 Post-sequencing processing and fungal identification

We used the DADA2 pipeline (Callahan *et al*. 2016) in R version 4.1 (R Core Team 2020) to remove primers, clean the sequences, and produce amplicon sequence variants (ASVs). ASVs were chosen over clustering sequences into operational taxonomic units (OTUs), because appropriate similarity cut offs differ for different fungal taxa, and ASVs are less biased (Callahan *et al*. 2017) and have been shown to provide better resolution for zeta diversity analyses (Bay *et al*. 2020). Primers were removed using “cutadapt” version 1.18 (Martin 2011) with up to 20% mismatches allowed, after which sequences were filtered to remove ambiguous bases using the “filterAndTrim” function. Paired-end reads were merged and chimeras removed. ASVs were then putatively named using the “assignTaxonomy” function with the UNITE database version 8.3 general release FASTA file (Kõljalg *et al*. 2020; Nilsson *et al*. 2018) as the reference database. This UNITE release uses a dynamic clustering threshold to create “species hypotheses” (Kõljalg *et al*. 2020; Nilsson *et al*. 2018), and species hypotheses represented by single sequences were included. We modified a small number of entries in the UNITE database to reflect recent taxonomic updates of particular *Cortinarius* species by Nilsen *et al*. (2020) and Nilsen *et al*. (2021) (Supporting Information Table S1). The “assignTaxonomy” function uses the Ribosomal Database Project Naïve Bayesian Classifier algorithm (Wang *et al*. 2007) to assign taxonomic levels from the reference database to the sample ASVs, using k-mers of size 8 and 100 bootstrap replicates. We used the default minimum bootstrap confidence level of 50 for assigning a taxonomic level.

Once putatively named, ASVs were classified into functional guilds using the FunGuild database version 1.1 (Nguyen *et al*. 2016). ASVs with a “highly probable” or “probable” confidence ranking of belonging to ectomycorrhizal species were retained for further analyses. The genus *Lactifluus* was also retained due to its known ectomycorrhizal status (De Crop 2016). All other ASVs were excluded from further analyses. Prior to analysis, we used the “cd-hit-est” server (Huang *et al*. 2010) to cluster all sequences at 100% similarity to merge any ASVs of different lengths that were otherwise identical. This merged 36 pairs of ASVs, for which the reads were summed and the longest sequence retained as the representative sequence.

After processing, 39,478 distinct ASVs were identified, of which 5,019 (17.7%) were classified as ectomycorrhizal. Ectomycorrhizal ASVs made up 58.7% of the sequence reads (7,943,552 of 13,522,723 total reads). This confirmed that the hyphal ingrowth bags were selectively filtering for ectomycorrhizal hyphae, because although ectomycorrhizal ASVs only accounted for a small proportion of total ASVs, they represented the majority of sequence reads. Read depth ranged between 539 and 47,443 per sample, and reads of ectomycorrhizal sequences ranged between 32 and 40,547, except for one sample which contained zero ectomycorrhizal reads, and was therefore excluded from further analysis. Despite the wide range in read depths, rarefaction curves of ectomycorrhizal sequences showed that all 807 samples with ectomycorrhizal reads were saturated, indicating that the read depth was sufficient to detect the vast majority of ASVs in each sample (Supporting Information Figures S1 and S2). Therefore, raw read counts were converted to presence/absence for each sample. We summarised the data for each of the 81 sites by using the number of plots each ASV was recorded in at that site as a measure of relative abundance. Therefore, each ASV received a value between 0 and 10 for each site.

### 3.5 Environmental sampling

Variables relating to host species, soil physical and chemical properties, forest structure, ground covers and topography were collected from each of the 81 survey sites. Additionally, we extracted points from GIS layers relating to climate, elevation, and solar radiation for each of the site locations (Table 2).

**Table 2:**
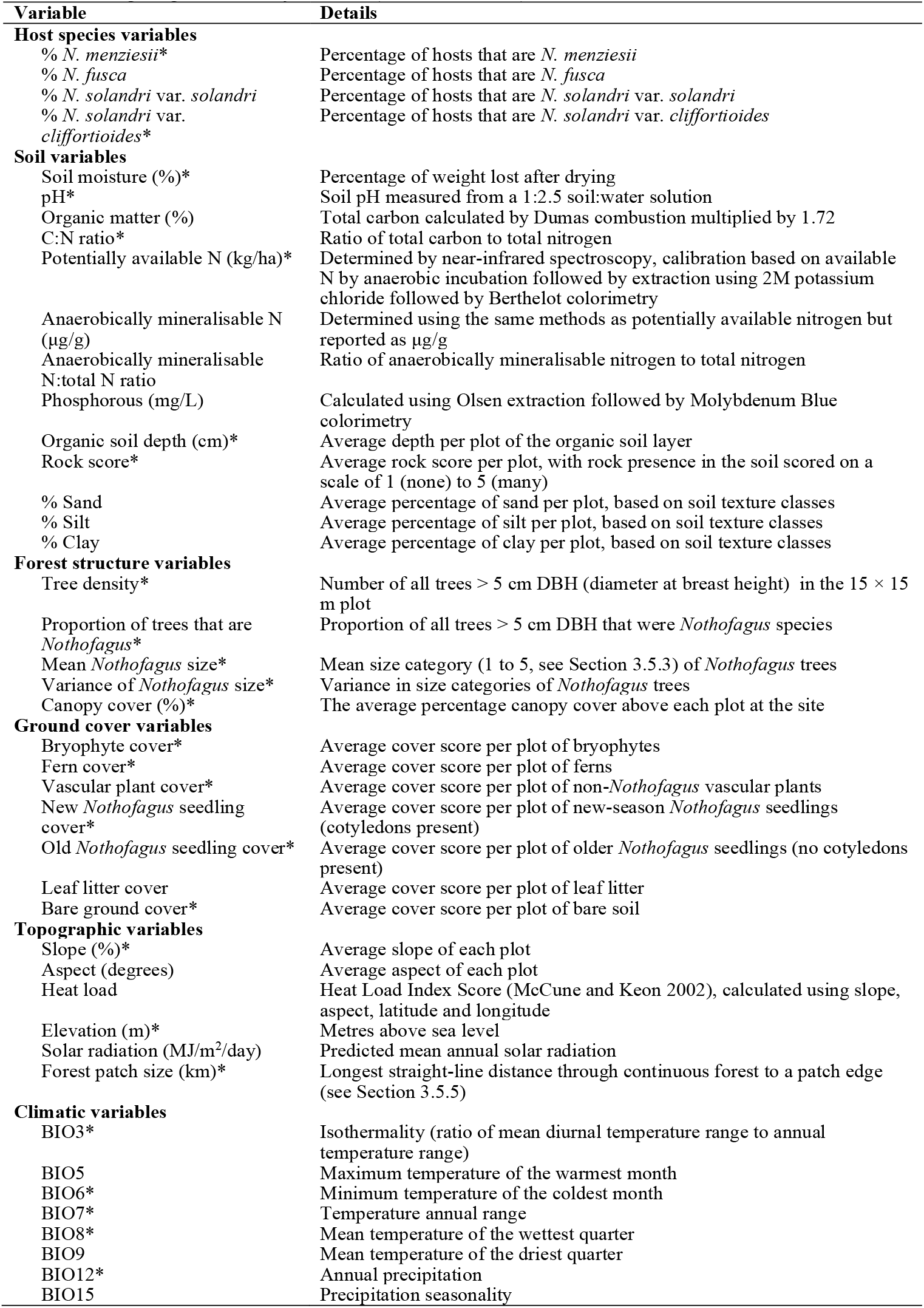
The environmental variables used in analyses. The maximum correlation between variables was r = |0.74| (Supporting Information Figures S3 and S4). * denotes the 27 variables used in the combined model assessing fungal community turnover (see Section 3.6.2).

#### 3.5.1 Host species

The host species at each site were characterised by estimating the percentage of *Nothofagus* trees belonging to a) *N. menziesii*, b) *N. fusca*, c) *N. solandri* var. *solandri*, and d) *N. solandri* var. *cliffortioides. Nothofagus truncata* only occurred at two sites, so it was not statistically possible to include the percentage of this species as a host variable.

#### 3.5.2 Soil properties

At the time of ingrowth bag collection, 3 soil cores (1.5 cm diameter, 10 cm deep) were taken from a random location within each of the 10 plots at each site (30 cores per site). Where possible, cores were positioned to capture the organic and mineral soil interface where the bags were buried (i.e., when the organic soil layer was deep, some organic matter was removed before taking the core). Cores from all 10 plots were mixed to form a single soil sample from each site. Samples were frozen in the field with dry ice and stored at −20 °C until analysed. Equipment was sterilised between plots with a 0.4% bleach solution.

For soil chemistry analysis, samples were de-frosted and homogenised. To calculate soil moisture content, 5 g sub-samples were weighed, dried overnight at 105 °C, and re-weighed. The remaining sample was dried at 38 °C, crushed, and sieved through a 1 mm sieve. We measured soil pH of 1:2.5 soil:water solutions. Additionally, organic matter content, carbon to nitrogen ratio, potentially available nitrogen, anaerobically mineralisable nitrogen, anaerobically mineralisable nitrogen to total nitrogen ratio, and Olsen phosphorous were measured by Hill Laboratories (https://www.hill-laboratories.com). Soil weight to volume ratio, total carbon, and total nitrogen were also measured by Hill Laboratories, but these variables were excluded from analyses due to being highly correlated (r > 0.82) with other soil variables. Soil moisture data was unavailable for one site, so the average of values was assigned to that site.

In addition to soil chemistry data, for each of the 10 plots we recorded the depth of the organic soil layer, scored the amount of rocks on a scale of 1 to 5, and classified the soil texture using the USDA soil texture classification classes (Soil Science Division Staff 2017). Soil texture classes were converted into the percentage of sand, silt and clay using the midpoints of each texture category on the USDA soil textural triangle (Soil Science Division Staff 2017). Plot values for organic soil layer depth, rock score, and percentage sand, silt and clay were averaged to provide a single value per site.

#### 3.5.3 Forest structure

To assess the forest structure of each site, we recorded the number of trees in five DBH (diameter at breast height) size categories in a 15 × 15 m plot situated in the middle of each site: 1) 5–10 cm, 2) 10–20 cm, 3) 20–50 cm, 4) 50–100 cm, and 5) > 100 cm. These data were used to calculate tree density, the mean and variance of *Nothofagus* stem sizes, and the proportion of trees that were *Nothofagus*. Mean and variance variables were calculated as the mean and variance of size category numbers (1 to 5) instead of centimetres because actual DBH measurements were not recorded for each tree. We also assessed canopy cover above each plot using the “%cover” canopy cover assessment app (https://percentagecover.com), and then averaged the covers to obtain a single value per site.

#### 3.5.4 Ground covers

To assess ground cover, we scored the percentage of each of the 10 plots at each site covered by bryophytes, ferns, non-*Nothofagus* vascular plants, new-season *Nothofagus* seedlings (cotyledons present), older *Nothofagus* seedlings (no cotyledons), leaf litter, and bare ground using Braun-Blanquet cover scores (0 = 0% cover, 1 = <1%, 2 = 1–5%, 3 = 6–25%, 4 = 26– 50%, 5 = 51–75%, and 6 = 76–100%). Plot cover scores (0 to 6) for each variable were averaged for each site.

#### 3.5.5 Topography

We recorded the slope and aspect of each plot and averaged the values for each site. The averaged slope and aspect, along with the latitude and longitude of the site, were used to calculate Heat Load Index scores (McCune and Keon 2002) in R (R Core Team 2020) using the “htld” function from the “ecole” package (Smith 2021). We also extracted elevation and mean annual solar radiation data for each site using the New Zealand Digital Elevation Model (Landcare Research 2010) and Mean Annual Solar Radiation (Landcare Research 2003) GIS layers. To calculate a measure of forest patch size, we recorded the furthest straight-line distance from the site location to a forest patch edge without intersecting non-forest areas on the EcoSat Forest mapping layer (Landcare Research 2014). This method was chosen to provide an indication of relative patch size rather than using patch area due to issues with calculating area for long continuous inter-connected patches.

#### 3.5.6 Climate

We downloaded point values for each site location of the 19 derived BIOCLIM variables relating to temperature and rainfall from the WorldClim dataset (Fick and Hijmans 2017) using the “raster” package (Hijmans 2020) in R (R Core Team 2020). Highly correlated variables were removed, resulting in eight of the 19 variables selected for analysis (Table 2) with correlations r < |0.63| (Supporting Information Figure S3).

### 3.6 Statistical analysis

All statistical analyses were conducted in R version 4.1 (R Core Team 2020). MS-GDM models were run using the New Zealand eScience Infrastructure (NeSI) high performance computing facilities (https://www.nesi.org.nz).

#### 3.6.1 Community assembly processes

To assess the influence of stochastic and deterministic processes on community assembly, we examined the rate and form of zeta diversity decline with increasing zeta order, along with orthodromic distance-decay relationships, using the “Zeta.decline.mc” function and “Zeta.ddecay” function from the “zetadiv” R package (Latombe *et al*. 2018a; McGeoch *et al*. 2019). Up to 10,000 random selections of site combinations were used, and regression lines for distance-decay were fitted using general additive models (GAMs). Sharp rates of decline in zeta diversity across the first few zeta orders indicate that turnover is mostly structured by differences in the composition of rare species (Latombe *et al*. 2019; McGeoch *et al*. 2019). The zeta ratio, or retention rate (zeta diversity of order *n* divided by zeta diversity of order *n*-1), provides a more precise method of visualising the tail of zeta decline. This describes the rate at which shared species are retained when an additional site is added to the comparison (i.e., the rate at which common species are retained across the landscape). If the retention rate declines towards zero after a particular zeta order, this indicates complete turnover in community composition, whereas a plateauing retention rate indicates there are widespread species shared across many sites (Latombe *et al*. 2019; McGeoch *et al*. 2019). The parametric form of zeta decline provides information regarding the relative probability of the occurrence of species across sites, which indicates the degree to which community assembly is dominated by stochastic or deterministic processes (McGeoch *et al*. 2019). An exponential decline in zeta diversity indicates all species have equal chance of occurring at a given site (i.e., stochastic assembly), whereas a power-law form of decline indicates a non-random pattern of species co-occurrence (i.e., deterministic community assembly) (McGeoch *et al*. 2019).

#### 3.6.2 Environmental drivers of turnover

We conducted MS-GDM (multi-site generalised dissimilarity modelling) for zeta orders 2 through 8 to determine the influence of explanatory variables on fungal community turnover, using the “Zeta.msgdm” function from the “zetadiv” R package. This function first selects random samples without replacement of groups of *n* sites and calculates zeta diversity (number of shared species) for each random sample. The environmental explanatory variable values are then summarised for each random sample based on the difference in range of values present, to provide a single value for each sample (Latombe *et al*. 2019). Continuous variable values are summarised by first transforming the variable into isplines, and then averaging the difference of each ispline between all pairs of the *n* sites (Latombe *et al*. 2019). The average orthodromic distance between any two of the *n* sites is also calculated and transformed into isplines to provide an indication of the geographical proximity of the sites within the sample. The “Zeta.msgdm” function then uses generalised linear models to assess the relationship between zeta diversity and the new explanatory variable values for the random samples of sites selected.

We first ran six initial base MS-GDM models to investigate the separate effects of host species, soil variables, forest structure, ground covers, topography and climatic variables on species turnover. The most influential variables from the base models were then selected for inclusion in a combined model, to test the relative influence of important explanatory variables. For all models, three isplines and up to 10,000 random combinations of groups of *n* sites were used. For zeta orders where more than 10,000 combinations of groups were possible, we repeated the model 30 times and computed the average ispline values for each explanatory variable for the 30 replications. The ispline values indicate the amount of community turnover that occurs across each environmental gradient, so to select variables to include in the combined model, we extracted the maximum ispline score from the base models for each variable at each zeta order. For each zeta order, the maximum ispline scores from all base models were then standardised to range between 0 and 1, and we selected all explanatory variables with scores greater than 0.4. This allowed the inclusion of variables important at all zeta orders in the combined model. Twenty-seven explanatory variables met this criterion (Table 2). The 0.4 cut-off was chosen because this included the variables that appeared to capture the majority of community turnover across all base models. Isplines for each base model are provided in Supporting Information (Figures S5–S10).

The combined model was performed using the same parameters as the base models, with 30 replicates of the 10,000 random combinations of groups of *n* sites for each zeta order. We partitioned variance explained by host variables, all other environmental variables, and geographical distance using a modified version of the “Zeta.varpart” function (which currently only allows the partitioning of two groups). Variance partitioning of each of the 30 replicates was averaged for each zeta order.

To check that including 27 variables in a single model did not lead to overfitting, we used cross-validation (Refaeilzadeh *et al*. 2009) by splitting the 81 sites into training and testing data (75% and 25%, respectively). We built MS-GDM models with the training data for zeta orders 2 and 8 as described above, using up to 5,000 random combinations of groups of two and groups of eight sites. Cross-validation was not carried out for intermediate zeta orders due to computational constraints. Models were first run with two randomly selected explanatory variables, with additional variables added each time in a random order until all 27 variables were included. Models built with the training data were evaluated by using the “Predict.msgdm” function to predict the test data, and then calculating Pearson’s R^2^ between the predicted test data and actual test data values as a measure of variance explained. Training models were also used to predict the training data in the same way. Overfitting is evident if the variance explained when predicting the test data begins to decline as additional explanatory variables are included, when at the same time the variance explained when predicting the training data continues to increase. Each model was repeated at least 100 times with different random selections of training and testing sites and variables added in different random orders, to calculate the mean and standard error of the variance explained with increasing number of explanatory variables. The variance explained had not begun to decline when all 27 variables were included (Supporting Information Figure S11), showing that overfitting was not occurring.

To compare the relative influence of different types of environmental variables on ectomycorrhizal fungal community turnover, we calculated the variance explained by each of the six initial base models and the combined model of top variables for each of the zeta orders. To do this we used the same cross-validation approach described above by splitting the 81 sites into training data (75%) and testing data (25%). We built training models with up to 5,000 random combinations of *n* sites for each zeta order, and used these to calculate the variance explained when predicting the test data. Each model was repeated at least 100 times with different random selections of training and testing sites.

#### 3.6.3 Characterising communities

To assist with interpreting patterns in turnover, we calculated the proportion of ASVs at each site that belonged to the seven most diverse genera: *Cortinarius, Ruhlandiella, Inocybe, Tomentella, Laccaria, Russula*, and *Sebacina*. We ran general additive models (GAMs) on log-transformed proportion data using the environmental variables most strongly linked to turnover patterns, to examine how the relative diversity of different genera changed along the environmental gradients. GAMs were conducted using the “gam” function from the “mgcv” R package (Wood 2011) using Gaussian distributions, REML as the smoothing parameter estimation method, thin plate regression splines, and seven basis functions (k). The “gam.check” function was used to produce diagnostic plots for each model to ensure accurate model fit, and to check that the number of basis functions was not too low.

Additionally, we classified ASVs belonging to different genera into hyphal exploration type categories, to examine how the environmental variables affect ectomycorrhizal functional types. Ectomycorrhizal fungi explore the surrounding substrate using extramatrical mycelia, which can spread far or be concentrated close to the root, and have smooth or dense structure (Agerer 2001). Agerer (2001) classified different types of mycelium into “exploration types”, and Tedersoo and Smith (2013) compiled available information regarding the exploration types of different ectomycorrhizal genera. The exploration type has strong influence on the enzymatic activity of ectomycorrhizal root tips and therefore nutrient uptake (Tedersoo *et al*. 2012b). We used the information provided by Tedersoo and Smith (2013) to classify ASVs into exploration types based the genus they were identified as (Table 3 and Supporting Information Table S2). We then grouped exploration types into two categories, those with dense or long hyphae (long distance, medium distance mat and medium distance fringe exploration types) and those with smooth or short hyphae (contact, short distance, and medium distance smooth types). This division follows that used by Geml *et al*. (2017), and best reflects differences in nutrient uptake strategies (Hobbie and Agerer 2010; Tedersoo and Smith 2013). ASVs belonging to genera known to contain species with multiple exploration types from both categories were excluded, as well as those for which the exploration type is currently unknown. We ran GAMs to examine the relationship between the proportion of ASVs belonging to each exploration category and environmental variables using the same methods for the genera described above, although response variables were normally distributed and so untransformed.

**Table 3:**
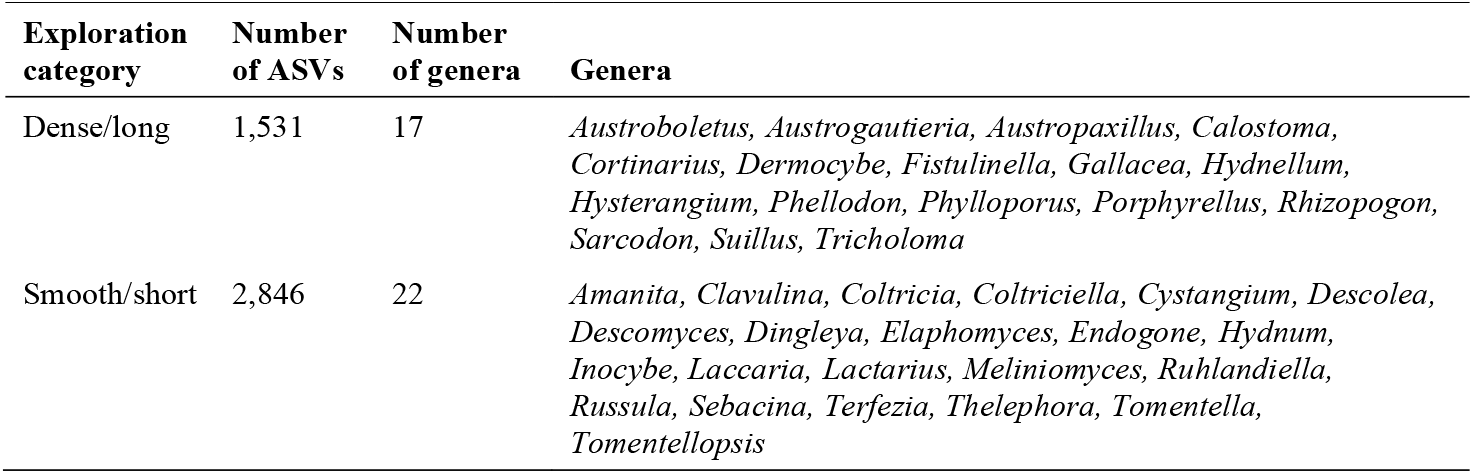
ASVs and genera classified as having “dense/long” or “smooth/short” hyphal exploration strategies. See Supporting Information (Table S2) for further classification details.

We also assessed ASV richness (alpha diversity) patterns. Because variables affecting total richness may be different to those influencing compositional turnover, we tested all environmental variables using GAMs. As with the MS-GDM analysis, we ran six initial base models for each variable category outlined in Table 2 (Supporting Information Figures S12– S17). All variables with *p*-values greater than 0.1 were then selected for inclusion in a combined model. Model parameters were as outlined above, with the number of basis functions (k) set to the highest value possible for each model whilst not exceeding the number of unique covariate combinations (k ranged between 5 and 17).

## 4. Results

A total of 5,019 ectomycorrhizal fungal ASVs comprising 7,943,552 sequence reads were identified across the 81 sites, with individual site richness ranging from 79 to 297. The majority of ectomycorrhizal ASVs belonged to the phyla Basidiomycota (81%) and Ascomycota (19%), and six ASVs from phylum Mucoromycota (genus *Endogone*) were also detected (Figure 2). Based on the putative taxonomic ASV identification, the most diverse genera present within the dataset were *Cortinarius* (28% of ASVs), *Ruhlandiella* (16%), *Inocybe* (11%), *Tomentella* (8%), *Laccaria* (5%), *Russula* (4%), and *Sebacina* (4%; Figure 2).

**Figure 2:**
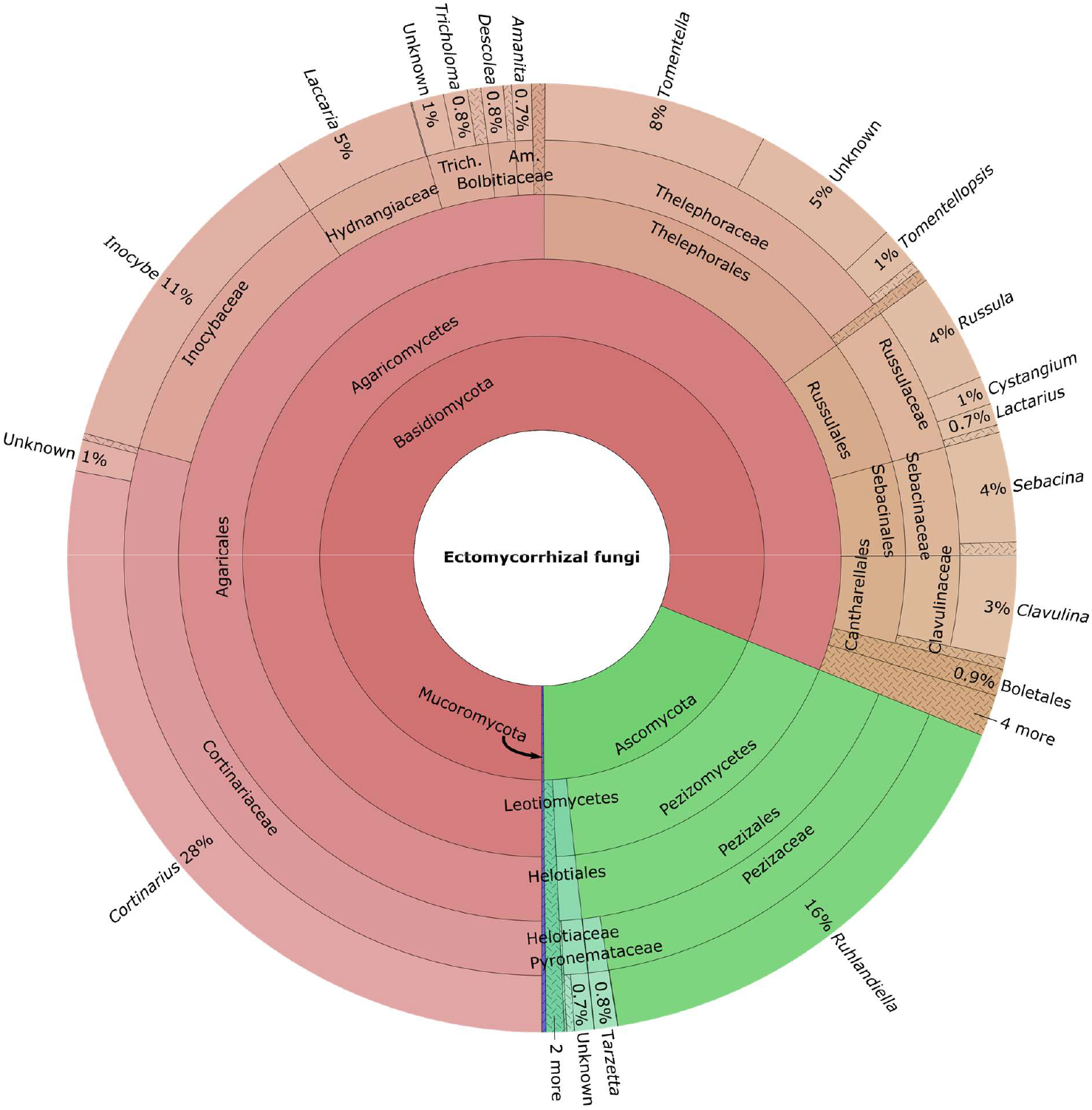
Krona chart showing the percentage of ASVs belonging to each taxonomic group, from phylum (inner circle) to genus (outer circle). Taxonomic names are those assigned from the UNITE database (see Section 3.4). Hashed segments indicate further divisions are present but not shown. Trich. = Tricholomataceae, Am. = Amanitaceae.

### 4.1 Community assembly processes

The zeta diversity analysis of the relative effect of deterministic and stochastic processes driving community assembly showed that the parametric form of decline in zeta diversity (shared ASVs) with increasing zeta order (number of sites included in the comparison) was clearly best explained by a power-law relationship rather than an exponential relationship (AIC = −16.4 and 15.8, respectively). This indicates that community assembly processes of this system are dominated by deterministic rather than stochastic processes, because there are uneven probabilities of the occurrence of ASVs across sites. Zeta diversity declined rapidly with increasing zeta order, with an average of less than two ASVs shared between groups of five sites or more and an average of less than one shared between more than eight sites (Figure 3a). This sharp decline highlights the strong influence of rare ASVs on community turnover. The retention rate (Figure 3b) plateaued at a zeta ratio of approximately 0.85, indicating that there are still a number of widespread ASVs common across the region.

**Figure 3:**
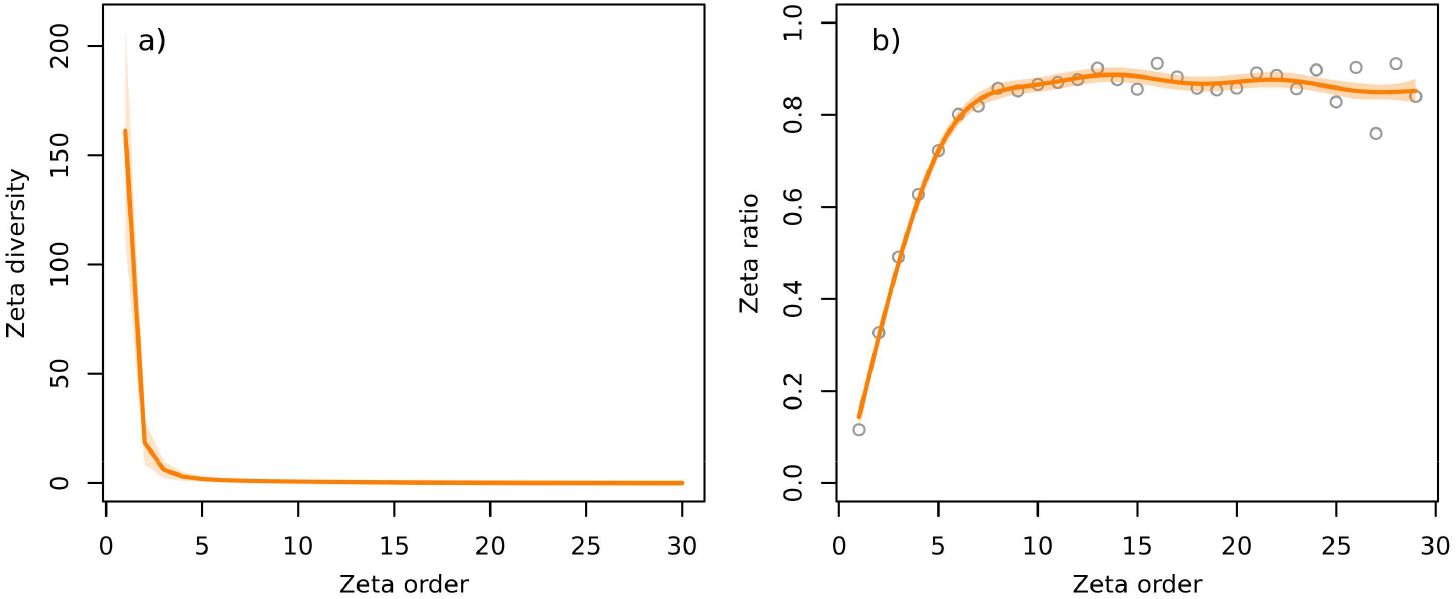
a) the decline in zeta diversity (number of shared ASVs) with increasing zeta order (number of sites included in the shared ASV calculations), and b) the zeta ratio (zeta diversity of order *n* divided by the zeta diversity of order *n*–1). The zeta ratio shows the retention rate, which is the proportion of shared species between a group of *n* sites that remain shared once a new site is added (McGeoch et al. 2019). The solid line shows a) the average zeta diversity of up to 10,000 different combinations of sites at each zeta order, and b) a smooth curve fitted using a GAM. Shading shows one standard deviation for a), and one standard error for b).

The limited effect of stochastic processes on community assembly was supported by the relationship between zeta diversity and distance between sites. There was a small decline in zeta diversity with increasing distance for zeta orders 2, 3 and 4 up to approximately 100 km, but the number of shared ASVs remained relatively constant for distances over 100 km (Figure 4). No relationships were observed between zeta diversity and distance for zeta orders greater than 5 (Figure 4), indicating that ASVs shared by groups of five or more sites are purely influenced by deterministic processes and not stochasticity, highlighting that environmental factors are likely driving patterns of turnover at this scale.

**Figure 4:**
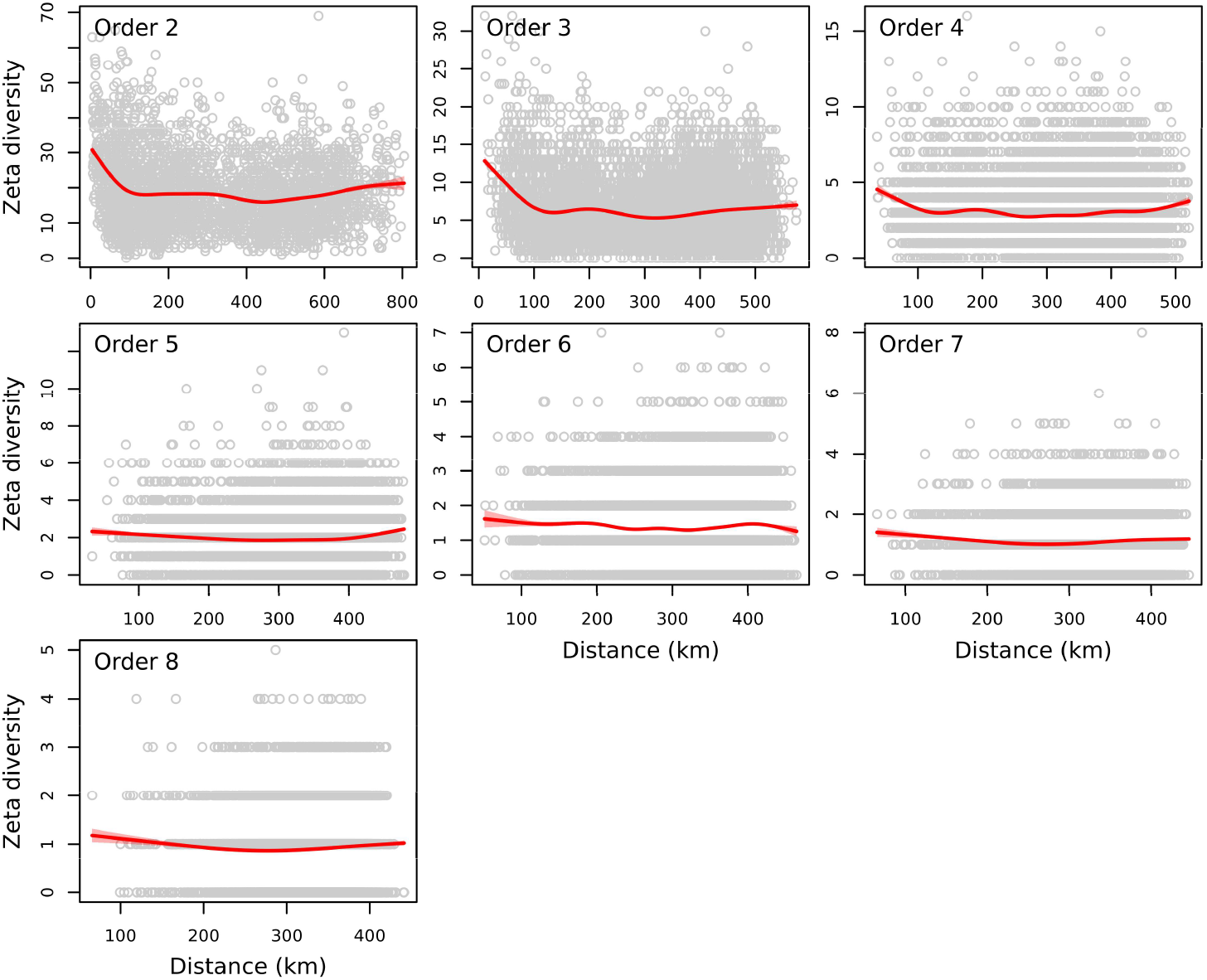
Changes in zeta diversity (number of shared ASVs) with increasing distance between sites. Points are 10,000 replications of randomly selected groups of *n* sites, and distances are averages of all pairs in the group. The solid red line shows a smooth curve fitted by a GAM, and pink shading shows one standard error.

### 4.2 Environmental drivers of turnover

Of the six base models testing the effect of different types of environmental variables on community turnover, ground cover variables had the strongest effect on turnover of rare ASVs (lower zeta orders), and soil variables most strongly affected turnover of more common ASVs (higher zeta orders; Figure 5). The host species model also performed well at lower zeta orders, but explained less variation than topography and forest structure at higher zeta orders (Figure 5). The climate model explained the least amount of variation at all zeta orders (Figure 5).

**Figure 5:**
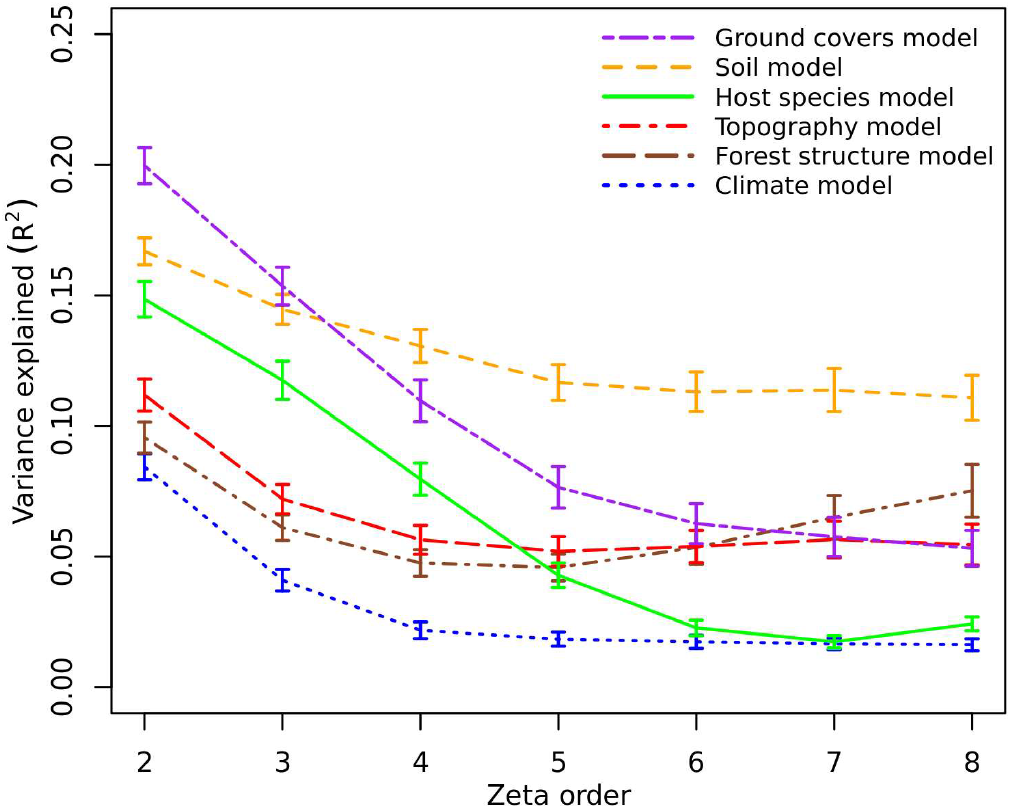
The variance explained (Pearson’s R^2^ calculated using cross validation) of the base models testing the effects of different types of environmental variables on fungal community turnover at different zeta orders, from zeta order 2 (pairs of sites) through to zeta order 8 (groups of eight sites). The variables included in each model are listed in Table 2. Lines are means and error bars one standard error of the iterations (200 for the soil model, 100 for other models; see Section 3.6.2).

The combined model (testing the relative effects of the 27 most influential variables from the base models) showed important effects of variables from all six categories on ectomycorrhizal turnover, but there were differences between those affecting rare and common ASV turnover (Figure 6). Soil C:N ratio and pH had relatively strong effects at all zeta orders (Figure 6). Vascular plant cover, bryophyte cover, organic soil depth, forest patch size and annual precipitation (BIO12) were strong drivers of turnover of rare ASVs, but not common ASVs. Conversely, temperature annual range (BIO7), isothermality (BIO3), mean *Nothofagus* size, and slope were drivers of common ASV turnover, but had less effect on rare ASVs (Figure 6). *Nothofagus* host species variables had moderate effects on rare ASV turnover, although these affects were less important than many other environmental variables (Figure 6). Variance partitioning confirmed that the majority of variance explained was directly attributed to environmental variables (28.5–35% for all zeta orders), but some variance could also only be explained by host variables, particularly at lower zeta orders (3.8% for zeta order 2; Figure 7). There was also substantial variance that could not be directly attributed to either host, environment, or distance effects (overlapping segments in Figure 7). It is therefore possible that host effects account for up to 13.7% of variance explained, although it is also possible some of this additional variance is the result of environment or distance effects. As indicated by the distance-decay relationships in Figure 4, distance between sites moderately influenced rare ASV turnover (Figure 6), but very little variance was attributed purely to distance (Figure 7), and distance effects were non-existent after zeta order 4.

**Figure 6:**
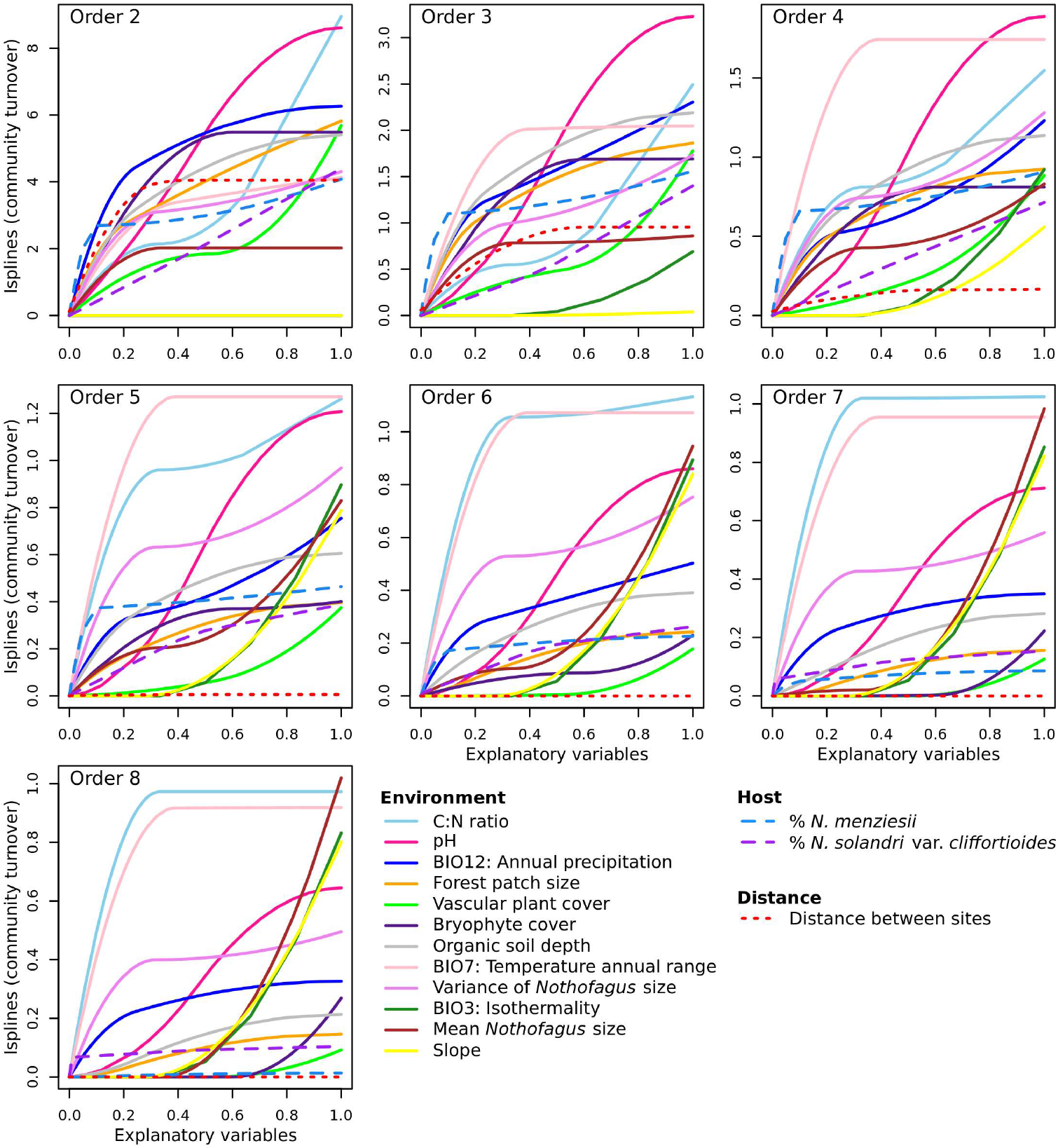
Isplines (average of 30 replicates) showing the contribution of environmental variables, host variables, and distance between sites to the combined model of top variables explaining zeta diversity at each zeta order. For the environment category, only variables ranked in the top six for at least one of the orders are shown (12 out of 25). Only two host variables met the criterion for inclusion in the combined model (see Table 2 and Section 3.6.2). The x-axes show the original predictor variable values rescaled between 0 and 1. The relative amplitude of each ispline in each panel indicates the relative importance of that variable for explaining zeta diversity, and therefore the effect of that variable on ectomycorrhizal community turnover. The slope of the ispline indicates the rate of community turnover occurring at different points along the environmental gradient. Isplines for all variables and the 30 replicates, along with original variable units, are provided in Supporting Information Figures S18–S24.

**Figure 7:**
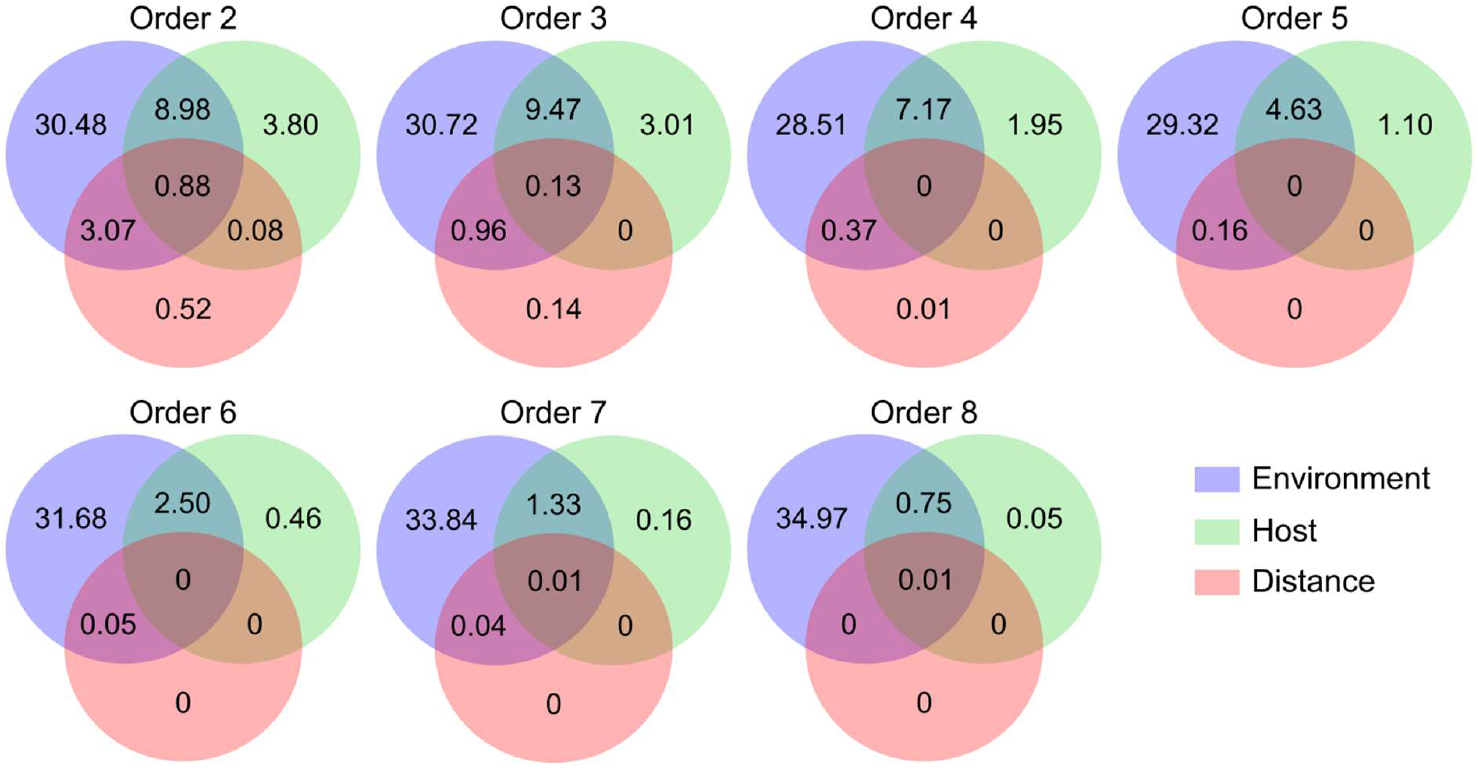
Variance partitioning of the different types of variables included in the combined model (25 environment variables, 2 host variables, and distance between sites). Values are the percentage of variance explained.

### 4.3 Characterising communities

The relative diversity of different genera varied along environmental gradients, particularly in relation to soil properties, annual precipitation (BIO12) and variance in *Nothofagus* size (Figure 8). Changes in the proportion of *Cortinarius* ASVs commonly showed opposite patterns to other genera (Figure 8). ASVs thought to exhibit different hyphal exploration strategies also responded differently to some environmental variables, particularly C:N ratio, cover of vascular plants, and isothermality (BIO3; Figure 9). Most notably, the proportion of ASVs with dense/long hyphal types was positively related to C:N ratio, whereas the proportion of ASVs with smooth/short types showed the opposite effect (Figure 9).

**Figure 8:**
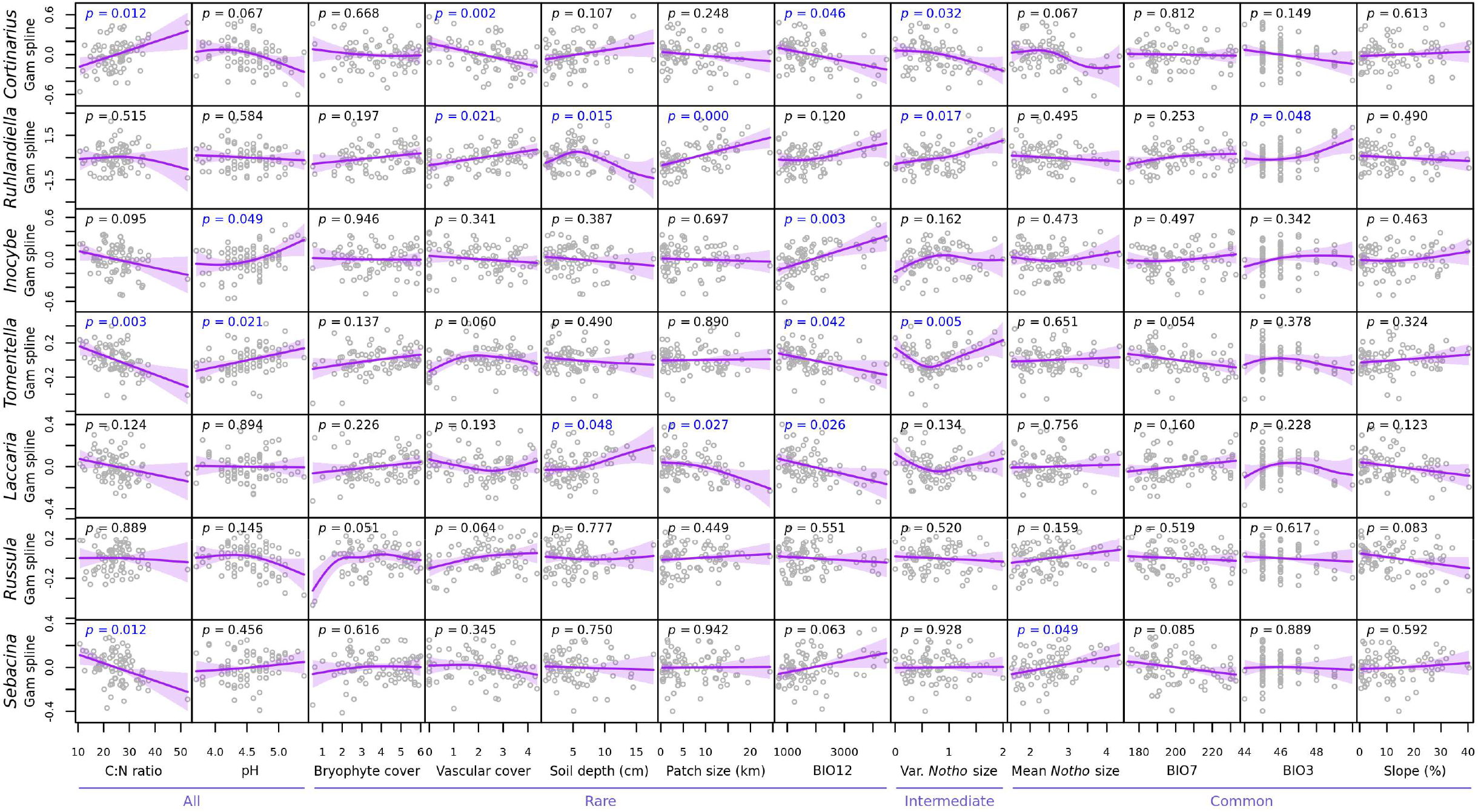
Partial residuals of fitted GAMs testing the effect of the top environmental variables presented in Figure 6 (x-axes) on the proportion of ASVs belonging to each of the seven most diverse genera (y-axes). Shading shows one standard error of the fitted smooth curves. Whether environmental variables predominantly influenced all, rare, intermediate, or common ASV turnover is indicated below variable names. Significant *p*-values (< 0.05) are in blue. BIO12 = Annual precipitation (mm), BIO7 = Temperature annual range (°C × 10), BIO3 = Isothermality. Var. *Notho* size = variance of *Nothofagus* size.

**Figure 9:**
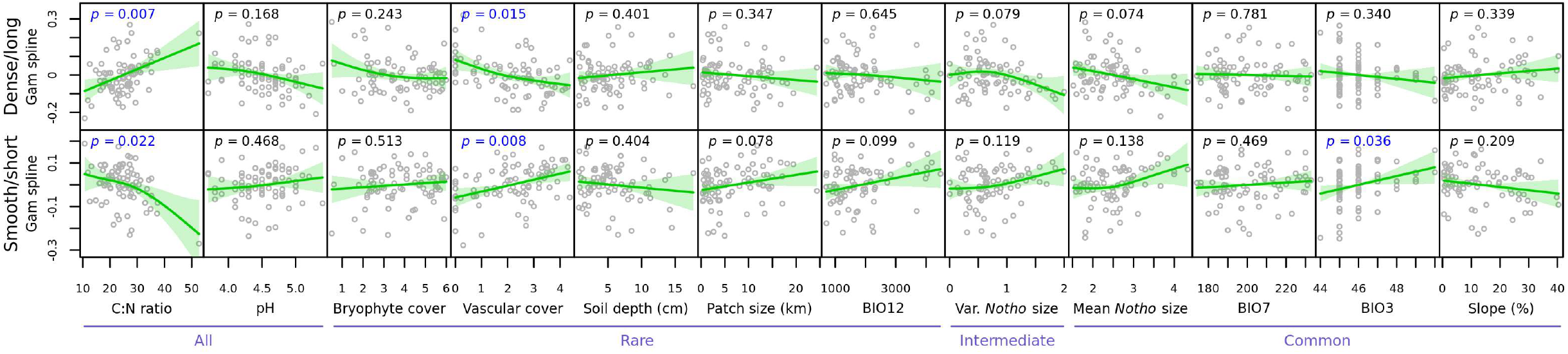
Partial residuals of fitted GAMs testing the effect of the top environmental variables presented in Figure 6 (x-axes) on the proportion of ASVs belonging to dense/long or smooth/short hyphal exploration types (y-axes). See Table 3 for details of the genera included in each exploration category. Shading shows one standard error of the fitted smooth curves. Whether environmental variables predominantly influenced all, rare, intermediate, or common ASV turnover is indicated below variable names. Significant *p*-values (< 0.05) are in blue. BIO12 = Annual precipitation (mm), BIO7 = Temperature annual range (°C × 10), BIO3 = Isothermality. Var. *Notho* size = variance of *Nothofagus* size.

Variation in ASV richness (alpha diversity) was best explained by elevation, soil rock score (negative relationships), and annual precipitation (positive relationship; Figure 10). Although bryophyte cover and soil moisture were significantly positively related to richness in the initial ground covers and soil models (Supporting Information Figures S13 and S15), these effects were absent from the combined model of best variables (Figure 10), likely due to variation being better explained by annual precipitation. No host variables were significantly related to ASV richness (Supporting Information Figure S12)

**Figure 10:**
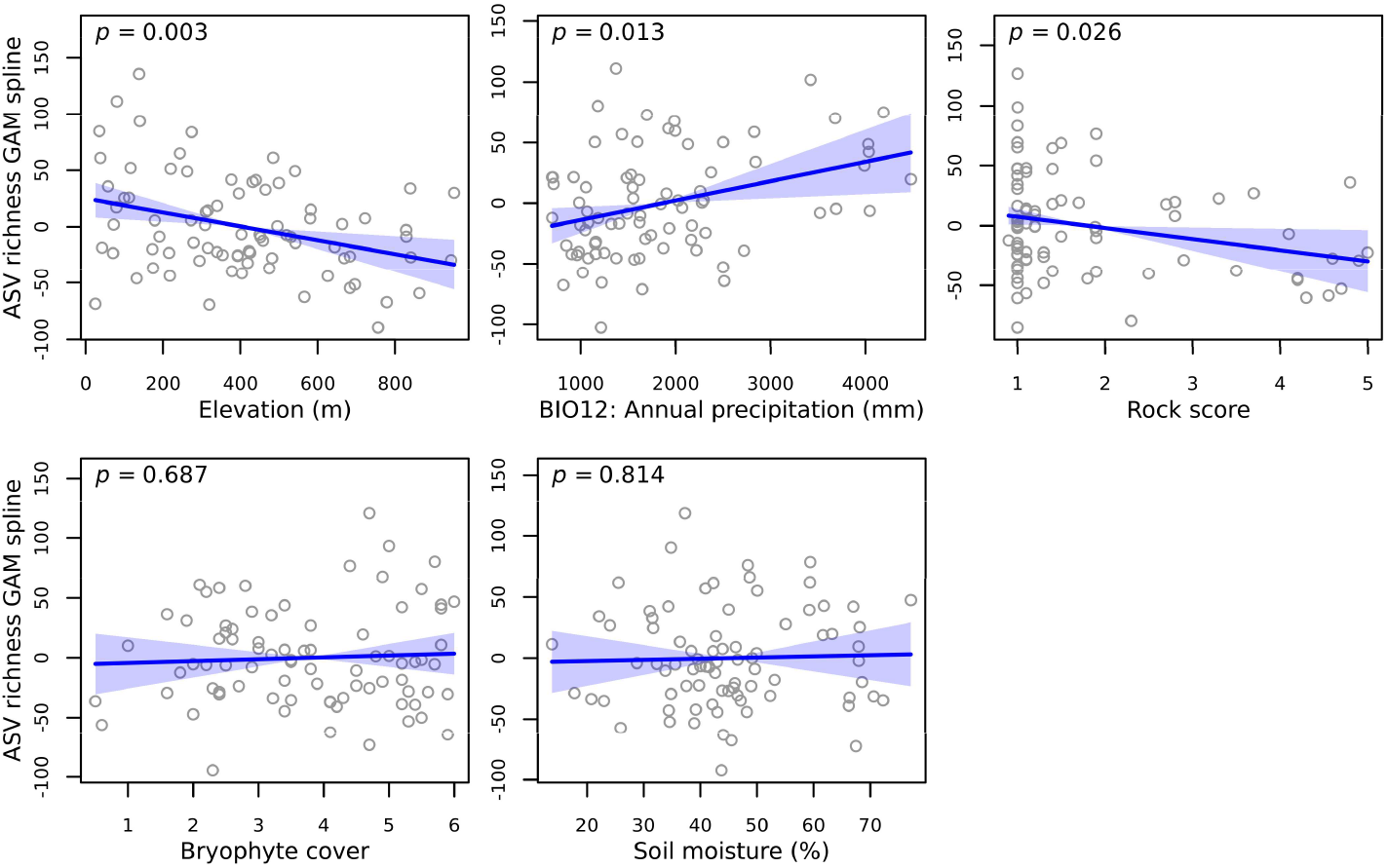
Partial residuals of the combined GAM model testing the relationship between important environmental variables from the initial base models (Supporting Information Figures S12–S17) and ASV richness. Deviance explained = 32.3%, adjusted R^2^ = 0.28. The solid line shows the fitted smooth curve and shading shows one standard error.

## 5. Discussion

By applying the zeta diversity analysis framework to ectomycorrhizal fungal communities, this study has shown that New Zealand ectomycorrhizal community assembly is primarily driven by deterministic selection, and that different environmental variables drive turnover of rare and common species. Soil C:N ratio and pH drove turnover patterns across the full range of rare, intermediate, and widespread ASVs. Ground cover variables, soil depth, annual precipitation and forest patch size had strong effects on rare taxa, whereas temperature variables, host tree size and slope had stronger effects on more widespread taxa. Some variance in turnover was also directly linked to host species identity, particularly at lower zeta orders (i.e., rare ASV turnover). Elevation had less important effects than many other variables on turnover, but was important for explaining alpha diversity patterns. These results confirm that pairwise comparisons of beta diversity reflect only part of community compositional turnover dynamics, and highlight the benefits of conducting more thorough analyses that take into account different aspects of the community and regional species pool.

### 5.1 Community assembly processes

The importance of deterministic assembly processes found here aligns with increasing evidence that ectomycorrhizal fungal communities are strongly affected by the environment (Tedersoo *et al*. 2012a; van der Linde *et al*. 2018). Deterministic processes have been shown to dominate soil fungal community assembly in many countries (Beck *et al*. 2015; Chen *et al*. 2018; Dumbrell *et al*. 2010; Glassman *et al*. 2017; Kranabetter *et al*. 2015; Marion *et al*. 2021; Taylor *et al*. 2014), although stochastic processes have also been identified as important to varying degrees in some systems (Bahram *et al*. 2013; Chen *et al*. 2018; Dumbrell *et al*. 2010; Glassman *et al*. 2017).

Distance-decay patterns in our data did show evidence of stochastic effects across spatial scales up to 100 km for lower zeta orders, indicating that processes such as dispersal, speciation and ecological drift have some influence over rarer species (McGeoch *et al*. 2019). Dispersal ranges of ectomycorrhizal species are highly variable, depending on species traits and local site factors such as air currents (Peay and Bruns 2014). Whilst the spores of some species can travel long distances by wind or animal dispersal (Horton 2017), evidence of others indicates the vast majority of spores fall within one metre of the fruiting body (Galante *et al*. 2011), and some do not fruit above ground (Smith and Read 2008). Experimental evidence supports the idea that dispersal limitation likely has some effect on ectomycorrhizal community composition (Peay and Bruns 2014).

Quantifying possible effects of speciation and ecological drift is more difficult. This study used ASVs defined by the ITS region, which provides relatively fine separation of most taxa beyond taxonomic species level based on small differences in base pairs (Callahan *et al*. 2017). Rare taxa could therefore be ASVs undergoing the beginnings of speciation or those with random mutations as well as true range-restricted species. Using ASVs with zeta diversity analysis has been shown to enhance the detection of patterns that are lost when clustering sequences into OTUs (Bay *et al*. 2020), but the ecological importance of small differences separating closely related ASVs is difficult to quantify. Examining patterns in phylogenetic turnover would provide more information, but this cannot currently be implemented within the MS-GDM framework.

### 5.2 Environmental drivers of turnover

The fact that soil variables, particularly pH and C:N ratio, had the strongest effects on ectomycorrhizal turnover and influenced the full spectrum of rare, intermediate and common ASVs reflects the well-known specificity that many ectomycorrhizal taxa have for particular soil conditions (Horton *et al*. 2013; Krishna Sundari and Adholeya 2003; Suz *et al*. 2014). C:N ratio had particularly strong effects. We observed that the relative diversity of *Tomentella* and *Sebacina* was negatively related to C:N ratio, whereas *Cortinarius* diversity was positively related. These patterns are similar to findings by Kranabetter *et al*. (2015) along soil fertility gradients in Canadian *Pseudotsuga menziesii* forests, and reflect known differences in species’ nitrogen uptake strategies (Kranabetter *et al*. 2015; Pellitier and Zak 2018). When inorganic nitrogen levels are low, *Cortinarius* species have the capacity to access organic soil nitrogen using oxidative enzymes (Bödeker *et al*. 2014; Pellitier and Zak 2018). Conversely, *Tomentella* species are specialised in the uptake of ammonium and better adapted to nitrogen-rich sites (Kranabetter *et al*. 2015). The positive relationship between C:N ratio and the proportion of ASVs thought to exhibit long distance or dense hyphal exploration strategies, and the negative relationship between C:N ratio and the proportion of short and smooth exploration types, also aligns with these patterns. Short or smooth hyphal types are more common in mineral soils, and longer more complex hyphal structures tend to be specialised in organic nitrogen uptake (Hobbie and Agerer 2010; Tedersoo and Smith 2013).

Variables describing small-scale variation within sites, such as bryophyte and vascular plant ground covers and the depth or the organic soil layer, had stronger effects on rare ASVs than common ASVs. Small-scale site factors such as bryophyte cover provide habitat heterogeneity that could increase available niche space and promote the establishment of species dispersing from other areas (Holt 2009). Bryophyte cover has been shown to both promote the growth of ectomycorrhizal plant host seedlings (Wang *et al*. 2016), and directly increase ectomycorrhiza abundance (Cappellazzi *et al*. 2007). Non-ectomycorrhizal vascular plants also change soil fungal communities by providing hosts for arbuscular mycorrhizal fungi that are not supported by *Nothofagus* (Smith and Read 2008), which could impact less well-established ectomycorrhizal species through competition. Although, ectomycorrhizal fungi have been shown to exclude the establishment of arbuscular mycorrhizae (Becklin *et al*. 2012; Knoblochová *et al*. 2017). It is therefore possible that the link between vascular plants and ectomycorrhizal turnover is indirect and driven by other factors altering the ectomycorrhizal community in a way that allows vascular plants and their arbuscular symbionts to establish (e.g., Frouz *et al*. 2019).

Forest patch size (measured as the straight-line distance to the furthest patch edge) and annual precipitation (BIO12) were also linked to rare ASV turnover, particularly at the lower end of the gradients as patches increased in size from 300 m to c. 5 km (maximum patch size = 27 km) and annual precipitation increased from 700 mm to c. 1,300 mm (maximum = 4,046 mm). Forest fragmentation can negatively affect mycorrhizal root colonisation, diversity, and community composition due to altered nutrient cycling, environmental conditions, or increased pathogens (Grilli *et al*. 2017; Sapsford *et al*. 2020). This study detected no effect of patch size on total richness, but patch size was positively related to the relative diversity of *Ruhlandiella* and negatively related to the relative diversity of *Laccaria. Laccaria* relative diversity was also negatively related to precipitation. *Laccaria* is a widespread genus of fungi containing species adapted to many different conditions and successional stages (Wilson *et al*. 2017), possibly providing an advantage in drier regions or smaller patches where conditions may be more variable. *Ruhlandiella* is a genus of truffle-like species, but it has not previously been detected in New Zealand and little is known about its ecology (Kraisitudomsook *et al*. 2019).

In contrast to the small-scale factors influencing rare ASVs, turnover of more common ASVs was affected by large-scale climatic variables relating to temperature. Past studies show that climatic variables effect ectomycorrhizal fungi at global and regional scales (Jarvis *et al*. 2013; Miyamoto *et al*. 2015; Tedersoo *et al*. 2014; van der Linde *et al*. 2018; Větrovský *et al*. 2019), although the relative importance of climate compared to soil and plant characteristics varies depending on the scale and system examined (Matsuoka *et al*. 2016; Põlme *et al*. 2013; Rincón *et al*. 2015). Interestingly, we observed that turnover across the annual temperature range gradient occurred primarily at the lower end when the range increased from 17.4 °C– 19.5 °C (maximum = 23.3 °C). Furthermore, isothermality (ratio of mean diurnal temperature range to annual temperature range) mostly affected turnover at higher isothermality values when mean diurnal range (measured as mean of monthly (max temp − min temp)) was relatively high compared to annual temperature range. This indicates that short-term variation (diurnal range) has stronger effects than long-term variation (annual range) for common ASVs. Diurnal temperature range is predicted to increase in New Zealand under future climate scenarios (Ministry for the Environment 2018). Whilst this may not impact rare species in this system, these changes could have important implications for widespread ectomycorrhizal taxa.

Mean *Nothofagus* size also had strong effects on common ASV turnover, with the majority of turnover occurring at the higher end of the gradient and therefore driven by the presence of very large trees. Tree size likely reflects time since large-scale disturbance in this system, because *Nothofagus* forests are subjected to infrequent disturbances like landslides or fire that are stand-replacing (Stewart 1986). Ectomycorrhizal communities are known to follow postdisturbance successional processes in many regions (Bonet *et al*. 2004; Twieg *et al*. 2007; Visser 1995). Alternatively, it is possible that trees of different ages support different ectomycorrhizal species (Reverchon *et al*. 2012), although there is little published research regarding the influence of host age on mycorrhizal communities beyond the seedling stage. Mean *Nothofagus* size was positively related to the relative diversity of *Sebacina*, but not closely linked to other genera or hyphal exploration strategies.

Although elevation only weakly or moderately influenced turnover patterns at all zeta orders, elevation was one of the strongest predictors of alpha diversity, with richness declining as elevation increased. Similar negative relationships have been identified in other regions (Bahram *et al*. 2012; Kernaghan and Harper 2001; Nouhra *et al*. 2012), although some studies report richness peaks at mid elevations (Geml *et al*. 2017; Miyamoto *et al*. 2014). Elevation is a broad proxy for changes in climate, soil, and sometimes host communities (Bahram *et al*. 2012), and the weak influence of elevation on turnover could be because other variables included in our models capture more direct effects. However, the strong effect of elevation on alpha diversity indicates that the other variables assessed do not capture all changes that occur along the elevational gradient. Annual precipitation was also a strong driver of richness patterns, with greater richness in wetter regions. Interestingly, this contradicts global ectomycorrhizal patterns and experimental studies that show negative relationships between precipitation and diversity (Hawkes *et al*. 2011; Tedersoo *et al*. 2012a). However, other studies involving a wider range of soil fungi types show positive relationships (McGuire *et al*. 2012), indicating patterns may vary regionally.

### 5.3 Host versus environment

Although *Nothofagus* host identity variables had less influence on turnover than numerous other environmental variables, we observed medium-strength effects of host species on rare ASV turnover. Variance partitioning also showed that up to 3.8% of variance explained could be directly attributed to host effects, and host effects could possibly account for up to 13.7% of variation when considering variance that could not be separated from host and environment or distance. Whilst this value is smaller than the 28.5–35% of variance directly attributed to other environmental variables, it still provides evidence that host species identity explains variation in ectomycorrhizal turnover independently of the 25 other environmental variables included in the model. It is well known that many ectomycorrhizal species show affinities to particular host families or genera (Carriconde *et al*. 2019; Newton and Haigh 1998; Tedersoo *et al*. 2008; Tedersoo *et al*. 2013; van der Linde *et al*. 2018), but there is less evidence that specificity to particular hosts within the same genus also occurs. Nouhra *et al*. (2013) did not detect differences in below-ground ectomycorrhizal communities of South American *Nothofagus* species, although some differences in the identity of fruiting bodies were observed between forests dominated by different *Nothofagus* species in a nearby location (Nouhra *et al*. 2012). Therefore, the results of this study provide new evidence that ectomycorrhizal fungi may show some degree of preference to congeneric *Nothofagus* hosts.

Host effects could also be underestimated in this study, because we used percentage estimates to assess host presences at each site, which contain a margin of error and do not necessarily reflect below-ground root-fungus associations. We were also unable to incorporate phylogenetic relationships between host species due to limitations of the MS-GDM method, which has been shown to explain variation in ectomycorrhizal richness and community composition in other regions (Ishida *et al*. 2007; Tedersoo *et al*. 2013). More accurate quantification of host tree occurrences, incorporating phylogenetic relationships between host species, or assessing morphological characteristics of hosts likely to influence fungal associations could result in some currently unexplained variation being further attributed to host effects.

## 6. Conclusions

Zeta-diversity analysis has shown the importance of deterministic niche-based processes driving ectomycorrhizal community assembly in New Zealand, and revealed interesting differences in the types of environmental variables influencing rare and common ectomycorrhizal fungal turnover. Our analysis confirms that investigating turnover using zeta diversity captures more information than pairwise comparisons of beta diversity that are heavily influenced by rare taxa (Latombe *et al*. 2017). In particular, the stronger effects of temperature variables identified when examining turnover at higher zeta orders would have been missed using traditional analysis techniques, which has important implications for understanding the susceptibility of ectomycorrhizal communities to climate change (Větrovský *et al*. 2019).

Other studies using zeta diversity analyses have also shown that different environmental factors drive turnover of rare and common species for birds in Australia (Latombe *et al*. 2017), ants on islands in the Atlantic Ocean (Latombe *et al*. 2019), and fleas on small mammals (Krasnov *et al*. 2020b). Although, Ascensão *et al*. (2020) found no differences in drivers of turnover at different zeta orders of birds in Spain and Portugal. Interestingly, as observed here for fungi, studies in other systems also found that precipitation primarily affected rare species turnover, whereas temperature variables had greater effects on common species turnover (Krasnov *et al*. 2020b; Latombe *et al*. 2017; Latombe *et al*. 2019). That such disparate groups of organisms (fungi, birds, ants and fleas) show similar responses to climatic variables is intriguing, and raises the question of whether this pattern could be general. However, fleas exhibited opposite patterns in some biogeographical realms, possibly due to different realms containing phylogenetically dissimilar species that differ in their environmental tolerances (Krasnov *et al*. 2020b). Wider research is required before general cross-taxonomic trends can be identified.

The way in which diverging responses in rare and common species will impact ecosystem function is difficult to elucidate. Studies of plant and animal communities show that whilst rare species often contribute unique functional traits that increase functional diversity (Jain *et al*. 2014; Leitão *et al*. 2016; Mouillot *et al*. 2013), common species can have greater influence over functional processes due to their higher abundance (Jain *et al*. 2014). Therefore, whilst changes in variables driving rare species turnover could alter ectomycorrhizal functional diversity, changes in variables driving common species turnover could have more widespread impacts on ecosystem function. However, although our understanding of ectomycorrhizal species’ functional traits is improving (Agerer 2001; Tedersoo and Smith 2013), many species remain undescribed and their functional roles unknown (Rinaldi *et al*. 2008), so future work is required to understand whether patterns observed in other systems also hold true for fungi.

