## Supporting Information for "Different factors drive community assembly of rare and common ectomycorrhizal fungi"

**Table S1:** The *Cortinarius* sequence names modified and added to the UNITE version 8.3 database based on recent taxonomic work by Nilsen *et al.* (2020) and Nilsen *et al.* (2021).

| **Edited sequence names** | | | **New sequences added** | |
|---|---|---|---|---|
| **GenBank accession** | **Old name** | **New name** | **GenBank accession** | **Name** |
| MH101526 | *Cortinarius* sp. | *C. viridipileatus* | MT921419 | *C. taylorianus* |
| MK358101 | *Cortinarius* sp. | *C. violaceocystidiatus* | MW187061 | *C. sarchinochrous* |
| JX178616 | *C. taylorianus* | *Cortinarius* sp. | MN492664 | *C. pisciodorus* |
| MW187059 | *Cortinarius* sp. | *C. purpureocapitatus* | KJ635224 | *C. medioscaurus* |
| KT334135 | *Cortinarius* sp. | *C. minorisporus* | MT363100 | *C. epiphaeus* |
| MN492651 | *Cortinarius* sp. | *C. diaphorus* | MG367632 | *C. atropileatus* |
| AY669625 | *Cortinarius* sp. | *C. cuphocyboides* | | |
| AY033116 | *C. subcastinellus* | *C. cesarioanus* | | |
| AY033112 | *C. wallacei* | *C. subcastinellus* | | |
| JX178609 | *Cortinarius* sp. | *C. beeverorum* | | |

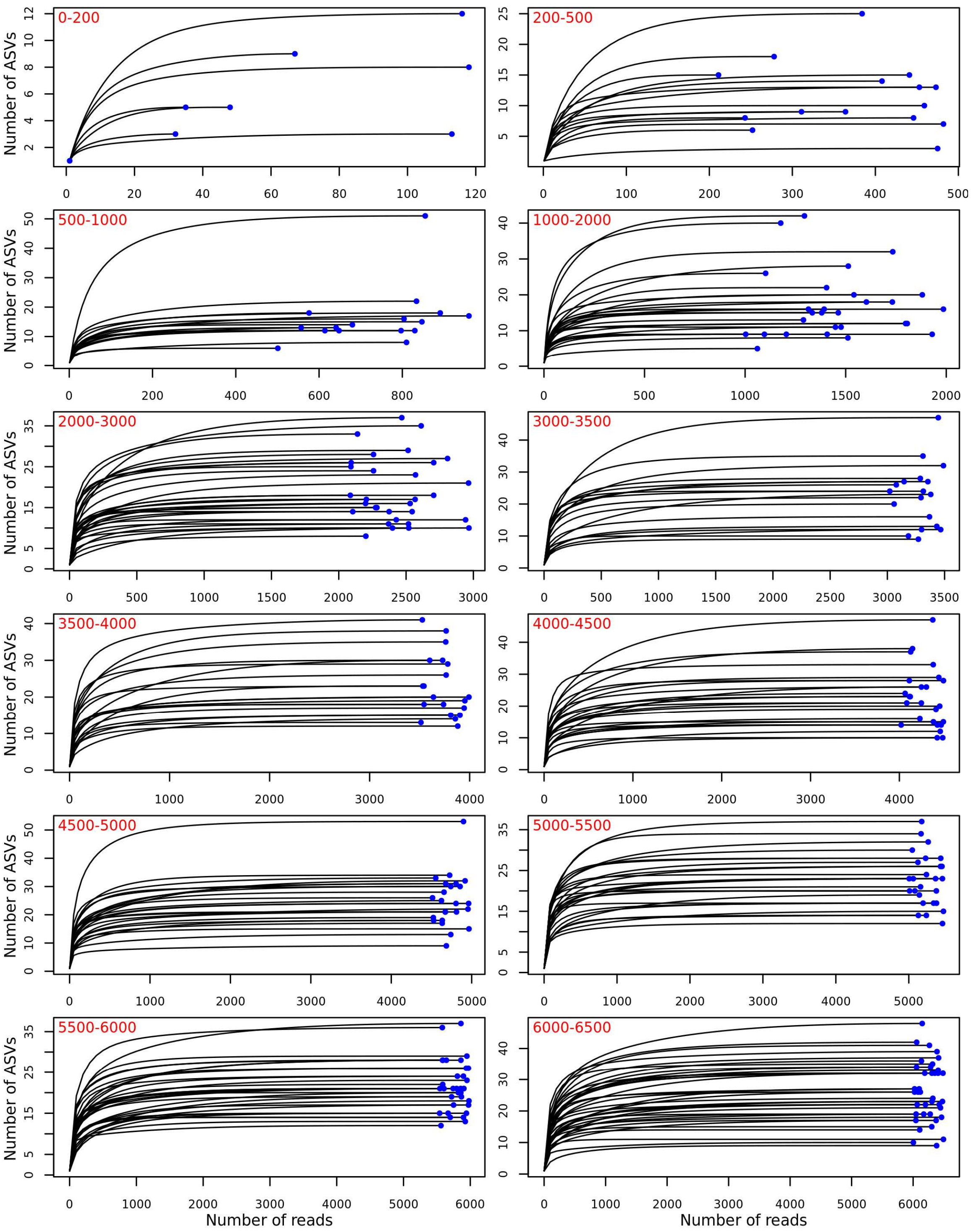


**Figure S1:** Rarefaction curves showing the increase in ectomycorrhizal ASVs detected (y-axis) with increasing read depth (x-axis). To assist with viewing, samples within particular final read depth ranges are plotted on different panels, with red text in the upper left of each plot indicating the range. Blue points show the final read depth for each sample. This figure contains panels for samples with read depths up to 6,500 reads. Samples with greater read depths are shown in Figure S2.

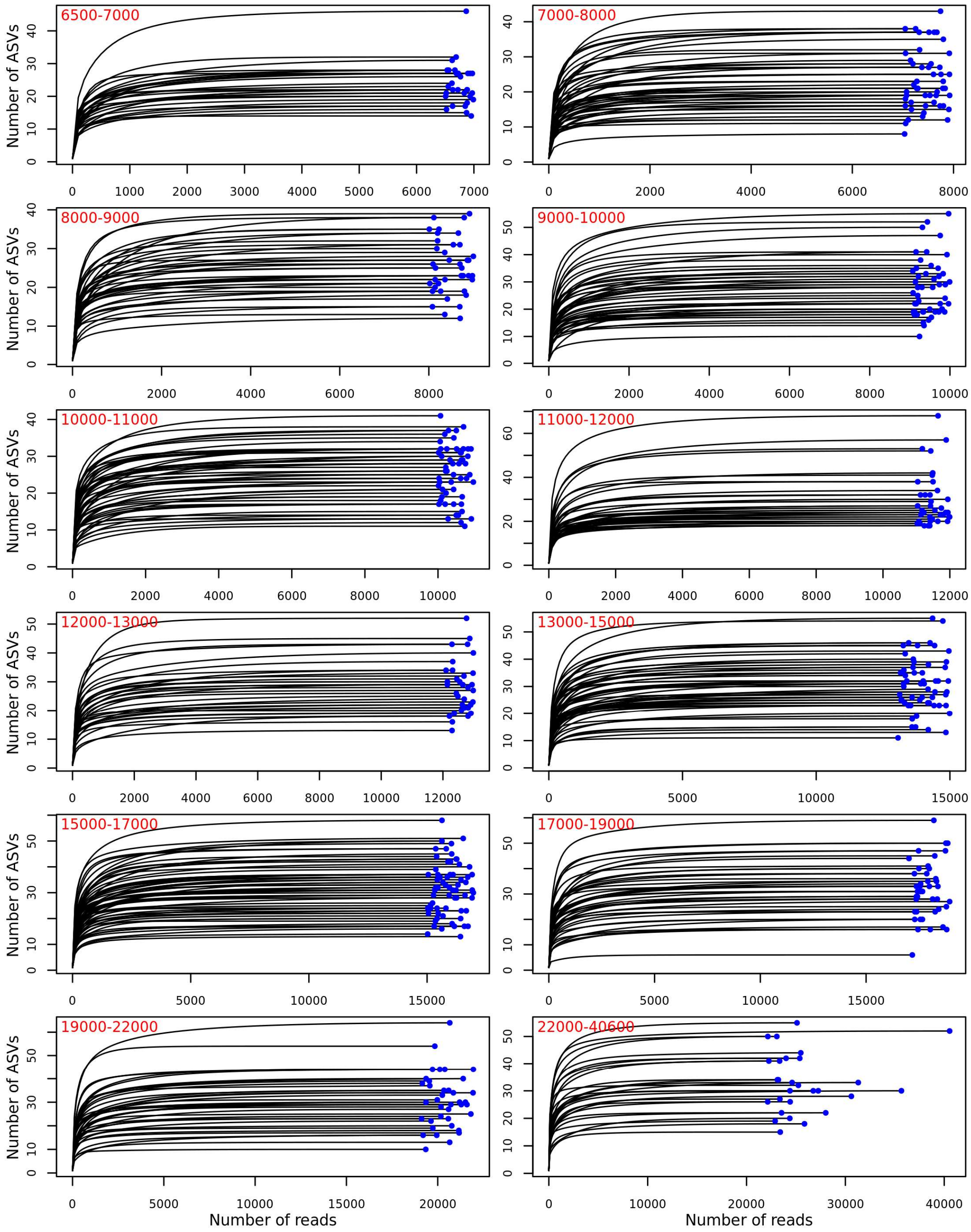


**Figure S2:** Rarefaction curves showing the increase in ectomycorrhizal ASVs detected (y-axis) with increasing read depth (x-axis). To assist with viewing, samples within particular final read depth ranges are plotted on different panels, with red text in the upper left of each plot indicating the range. Blue points show the final read depth for each sample. This figure contains panels for samples with read depths greater than 6,500 reads. Samples with fewer read depths are shown in Figure S1.

**a) Soil variables**

| | Organic soil depth | % Clay | % Silt | % Sand | Rock score | Soil moisture | pH | Phosphorous | P.A.N | A.M.N | A.M.N:Total N | Organic matter |
|---|---|---|---|---|---|---|---|---|---|---|---|---|
| Organic soil depth | 1.00 | | | | | | | | | | | |
| % Clay | 0.05 | 1.00 | | | | | | | | | | |
| % Silt | -0.04 | -0.24 | 1.00 | | | | | | | | | |
| % Sand | -0.10 | -0.53 | -0.17 | 1.00 | | | | | | | | |
| Rock score | -0.13 | -0.35 | 0.06 | -0.03 | 1.00 | | | | | | | |
| Soil moisture | 0.40 | -0.09 | 0.01 | -0.19 | 0.00 | 1.00 | | | | | | |
| pH | -0.42 | -0.25 | 0.12 | 0.25 | 0.07 | -0.28 | 1.00 | | | | | |
| Phosphorous | -0.10 | -0.21 | -0.14 | 0.15 | 0.15 | 0.23 | -0.07 | 1.00 | | | | |
| P.A.N | -0.37 | -0.09 | 0.05 | -0.15 | 0.09 | 0.04 | 0.53 | 0.07 | 1.00 | | | |
| A.M.N | -0.13 | -0.32 | -0.21 | -0.26 | 0.33 | 0.33 | 0.22 | 0.23 | 0.67 | 1.00 | | |
| A.M.N:Total N | -0.24 | 0.02 | -0.11 | -0.14 | -0.07 | -0.07 | 0.40 | -0.11 | 0.73 | 0.48 | 1.00 | |
| Organic matter | 0.22 | -0.28 | -0.28 | -0.21 | 0.35 | 0.48 | -0.22 | 0.25 | 0.17 | 0.74 | -0.06 | 1.00 |
| C:N ratio | 0.47 | 0.25 | -0.34 | -0.15 | -0.07 | -0.02 | -0.44 | -0.24 | -0.48 | -0.15 | -0.05 | 0.17 |

**b) Host species variables**

| | % *N. menziesii* | % *N. fusca* | % *N. solandri* var. *solandri* |
|---|---|---|---|
| % *N. menziesii* | 1.00 | | |
| % *N. fusca* | -0.22 | 1.00 | |
| % *N. solandri* var. *solandri* | -0.53 | -0.36 | 1.00 |
| % *N. solandri* var. *cliffortioides* | -0.30 | -0.19 | -0.26 |

**c) Ground cover variables**

| | Bryophyte cover | Fern cover | New *Notho* seedling cover | Old *Notho* seedling cover | Vascular cover | Leaf litter cover |
|---|---|---|---|---|---|---|
| Bryophyte cover | 1.00 | | | | | |
| Fern cover | -0.07 | 1.00 | | | | |
| New *Notho* seedling cover | 0.13 | -0.29 | 1.00 | | | |
| Old *Notho* seedling cover | 0.22 | -0.69 | 0.40 | 1.00 | | |
| Vascular cover | 0.07 | 0.52 | -0.44 | -0.58 | 1.00 | |
| Leaf litter cover | -0.74 | -0.09 | 0.12 | -0.06 | -0.25 | 1.00 |
| Bare ground cover | -0.32 | -0.16 | 0.11 | -0.07 | -0.15 | 0.20 |

**d) Forest structure variables**

| | Tree density | Tree prop. *Notho* | Mean *Notho* size | Variance *Notho* size |
|---|---|---|---|---|
| Tree density | 1.00 | | | |
| Tree prop. *Notho* | 0.06 | 1.00 | | |
| Mean *Notho* size | -0.42 | -0.70 | 1.00 | |
| Variance *Notho* size | -0.29 | 0.10 | 0.00 | 1.00 |
| Canopy cover | 0.05 | -0.28 | 0.06 | -0.13 |

**e) Climate variables**

| | BIO3 | BIO5 | BIO6 | BIO7 | BIO8 | BIO9 | BIO12 |
|---|---|---|---|---|---|---|---|
| BIO3 | 1.00 | | | | | | |
| BIO5 | -0.10 | 1.00 | | | | | |
| BIO6 | 0.05 | 0.28 | 1.00 | | | | |
| BIO7 | -0.12 | 0.57 | -0.63 | 1.00 | | | |
| BIO8 | 0.03 | 0.38 | 0.44 | -0.06 | 1.00 | | |
| BIO9 | -0.45 | -0.05 | 0.37 | -0.36 | -0.25 | 1.00 | |
| BIO12 | -0.35 | 0.23 | 0.24 | -0.02 | 0.09 | 0.02 | 1.00 |
| BIO15 | -0.29 | 0.11 | 0.33 | -0.19 | 0.13 | 0.26 | 0.31 |

**f) Topography variables**

| | Slope | Aspect | Heat load | Forest patch size | Elevation |
|---|---|---|---|---|---|
| Slope | 1.00 | | | | |
| Aspect | -0.10 | 1.00 | | | |
| Heat load | 0.56 | -0.55 | 1.00 | | |
| Forest patch size | -0.27 | 0.15 | -0.23 | 1.00 | |
| Elevation | 0.15 | -0.16 | 0.19 | -0.19 | 1.00 |
| Solar radiation | 0.16 | -0.09 | 0.19 | -0.01 | 0.56 |

**Figure S3:** Correlations (r) between the variables used in each base model analysis (see manuscript Table 2 for details). *Notho* = *Nothofagus,* P.A.N. = Potentially available N, A.M.N. = Anaerobically mineralisable N, Tree prop. *Notho* = Proportion of trees that are *Nothofagus*. See manuscript Table 2 for BIOCLIM variable details. Stronger correlations have darker shading.

### Combined model variables

| | Organic soil depth | Rock score | Soil moisture | pH | P.A.N | C:N ratio | BIO3 | BIO6 | BIO7 | BIO8 | BIO12 | Tree density | Tree prop. *Notho* | Mean *Notho* size | Variance *Notho* size | Canopy cover | Slope | Forest patch size | Elevation | Bryophyte cover | Fern cover | New *Notho* seedling cover | Old *Notho* seedling cover | Vascular cover | Bare ground cover | % *N. menziesii* |
|---|---|---|---|---|---|---|---|---|---|---|---|---|---|---|---|---|---|---|---|---|---|---|---|---|---|---|
| Organic soil depth | 1.00 | | | | | | | | | | | | | | | | | | | | | | | | | |
| Rock score | -0.13 | 1.00 | | | | | | | | | | | | | | | | | | | | | | | | |
| Soil moisture | 0.40 | 0.00 | 1.00 | | | | | | | | | | | | | | | | | | | | | | | |
| pH | -0.42 | 0.07 | -0.28 | 1.00 | | | | | | | | | | | | | | | | | | | | | | |
| P.A.N | -0.37 | 0.09 | 0.04 | 0.53 | 1.00 | | | | | | | | | | | | | | | | | | | | | |
| C:N ratio | 0.47 | -0.07 | -0.02 | -0.44 | -0.48 | 1.00 | | | | | | | | | | | | | | | | | | | | |
| BIO3 | 0.01 | -0.20 | -0.14 | 0.21 | 0.26 | 0.15 | 1.00 | | | | | | | | | | | | | | | | | | | |
| BIO6 | -0.14 | -0.14 | 0.23 | 0.11 | 0.39 | -0.22 | 0.05 | 1.00 | | | | | | | | | | | | | | | | | | |
| BIO7 | -0.12 | 0.11 | -0.21 | -0.09 | -0.32 | 0.17 | -0.12 | -0.63 | 1.00 | | | | | | | | | | | | | | | | | |
| BIO8 | -0.05 | -0.38 | 0.23 | -0.17 | 0.16 | 0.07 | 0.03 | 0.44 | -0.06 | 1.00 | | | | | | | | | | | | | | | | |
| BIO12 | 0.19 | -0.04 | 0.52 | -0.08 | -0.27 | -0.09 | -0.35 | 0.24 | -0.02 | 0.09 | 1.00 | | | | | | | | | | | | | | | |
| Tree density | -0.13 | 0.03 | -0.33 | -0.08 | -0.04 | 0.04 | -0.01 | -0.19 | 0.13 | -0.09 | -0.30 | 1.00 | | | | | | | | | | | | | | |
| Tree prop. *Notho* | 0.25 | -0.11 | -0.08 | -0.35 | -0.32 | 0.29 | -0.08 | -0.28 | 0.08 | -0.18 | -0.25 | 0.06 | 1.00 | | | | | | | | | | | | | |
| Mean *Notho* size | -0.12 | 0.16 | 0.26 | 0.16 | 0.07 | -0.05 | 0.07 | 0.16 | 0.00 | 0.16 | 0.42 | -0.42 | -0.70 | 1.00 | | | | | | | | | | | | |
| Variance *Notho* size | 0.12 | -0.09 | 0.21 | -0.07 | 0.05 | -0.01 | -0.10 | -0.02 | 0.18 | 0.07 | 0.15 | -0.29 | 0.10 | 0.00 | 1.00 | | | | | | | | | | | |
| Canopy cover | -0.08 | 0.13 | -0.03 | 0.15 | 0.17 | -0.17 | 0.05 | 0.39 | -0.30 | -0.10 | 0.19 | 0.05 | -0.28 | 0.06 | -0.13 | 1.00 | | | | | | | | | | |
| Slope | -0.22 | 0.45 | -0.23 | 0.16 | 0.17 | -0.12 | -0.10 | -0.20 | 0.18 | -0.21 | -0.30 | 0.11 | -0.20 | 0.07 | -0.05 | -0.07 | 1.00 | | | | | | | | | |
| Forest patch size | 0.25 | -0.12 | 0.18 | -0.20 | 0.00 | 0.14 | -0.02 | 0.40 | -0.26 | 0.19 | 0.24 | -0.11 | 0.06 | 0.01 | -0.02 | 0.08 | -0.27 | 1.00 | | | | | | | | |
| Elevation | 0.27 | 0.22 | 0.00 | -0.27 | -0.33 | 0.04 | -0.49 | -0.60 | 0.10 | -0.48 | -0.12 | 0.11 | 0.52 | -0.39 | 0.00 | -0.22 | 0.15 | -0.19 | 1.00 | | | | | | | |
| Bryophyte cover | 0.33 | -0.21 | 0.37 | -0.16 | -0.31 | 0.20 | 0.03 | 0.10 | -0.02 | 0.14 | 0.46 | -0.28 | 0.13 | 0.12 | 0.03 | -0.09 | -0.52 | 0.35 | -0.13 | 1.00 | | | | | | |
| Fern cover | -0.23 | -0.04 | 0.14 | 0.35 | 0.37 | -0.26 | 0.30 | 0.33 | -0.17 | 0.21 | 0.25 | -0.14 | -0.67 | 0.51 | 0.14 | 0.29 | -0.01 | 0.04 | -0.62 | -0.07 | 1.00 | | | | | |
| New *Notho* seedling cover | 0.06 | -0.04 | -0.10 | -0.13 | -0.19 | 0.37 | 0.32 | -0.21 | 0.40 | 0.04 | -0.29 | 0.16 | 0.18 | -0.10 | -0.09 | -0.25 | -0.05 | -0.15 | -0.10 | 0.13 | -0.29 | 1.00 | | | | |
| Old *Notho* seedling cover | 0.37 | 0.05 | -0.12 | -0.34 | -0.54 | 0.43 | -0.12 | -0.41 | 0.18 | -0.25 | -0.09 | 0.07 | 0.56 | -0.27 | -0.13 | -0.42 | -0.12 | -0.02 | 0.54 | 0.22 | -0.69 | 0.40 | 1.00 | | | |
| Vascular cover | -0.16 | -0.02 | 0.05 | 0.46 | 0.44 | -0.28 | 0.08 | 0.37 | -0.31 | 0.13 | 0.22 | -0.17 | -0.40 | 0.26 | 0.04 | 0.30 | 0.02 | 0.07 | -0.41 | 0.07 | 0.52 | -0.44 | -0.58 | 1.00 | | |
| Bare ground cover | -0.31 | 0.19 | -0.30 | 0.07 | -0.02 | -0.12 | -0.06 | -0.19 | 0.30 | -0.21 | -0.20 | 0.21 | -0.12 | 0.06 | -0.23 | -0.14 | 0.56 | -0.16 | 0.05 | -0.32 | -0.16 | 0.11 | -0.07 | -0.15 | 1.00 | |
| % *N. menziesii* | -0.18 | -0.30 | 0.23 | 0.13 | 0.21 | -0.19 | 0.08 | 0.34 | -0.17 | 0.40 | 0.29 | -0.18 | -0.47 | 0.40 | 0.05 | 0.20 | -0.31 | 0.10 | -0.54 | 0.18 | 0.59 | -0.24 | -0.58 | 0.44 | -0.20 | 1.00 |
| % *N. solandri* var. *cliffortioides* | -0.10 | -0.05 | -0.23 | 0.02 | -0.17 | -0.06 | -0.25 | -0.25 | 0.17 | -0.29 | -0.15 | 0.24 | 0.20 | -0.31 | -0.09 | -0.11 | 0.26 | -0.31 | 0.33 | -0.29 | -0.32 | 0.03 | 0.07 | -0.17 | 0.41 | -0.30 |

**Figure S4:** Correlations (r) between the variables used in the combined model (see manuscript Table 2 for details). *Notho* = *Nothofagus,* P.A.N. = Potentially available N, Tree prop. *Notho* = Proportion of trees that are *Nothofagus*. See manuscript Table 2 for BIOCLIM variable details. Stronger correlations have darker shading.

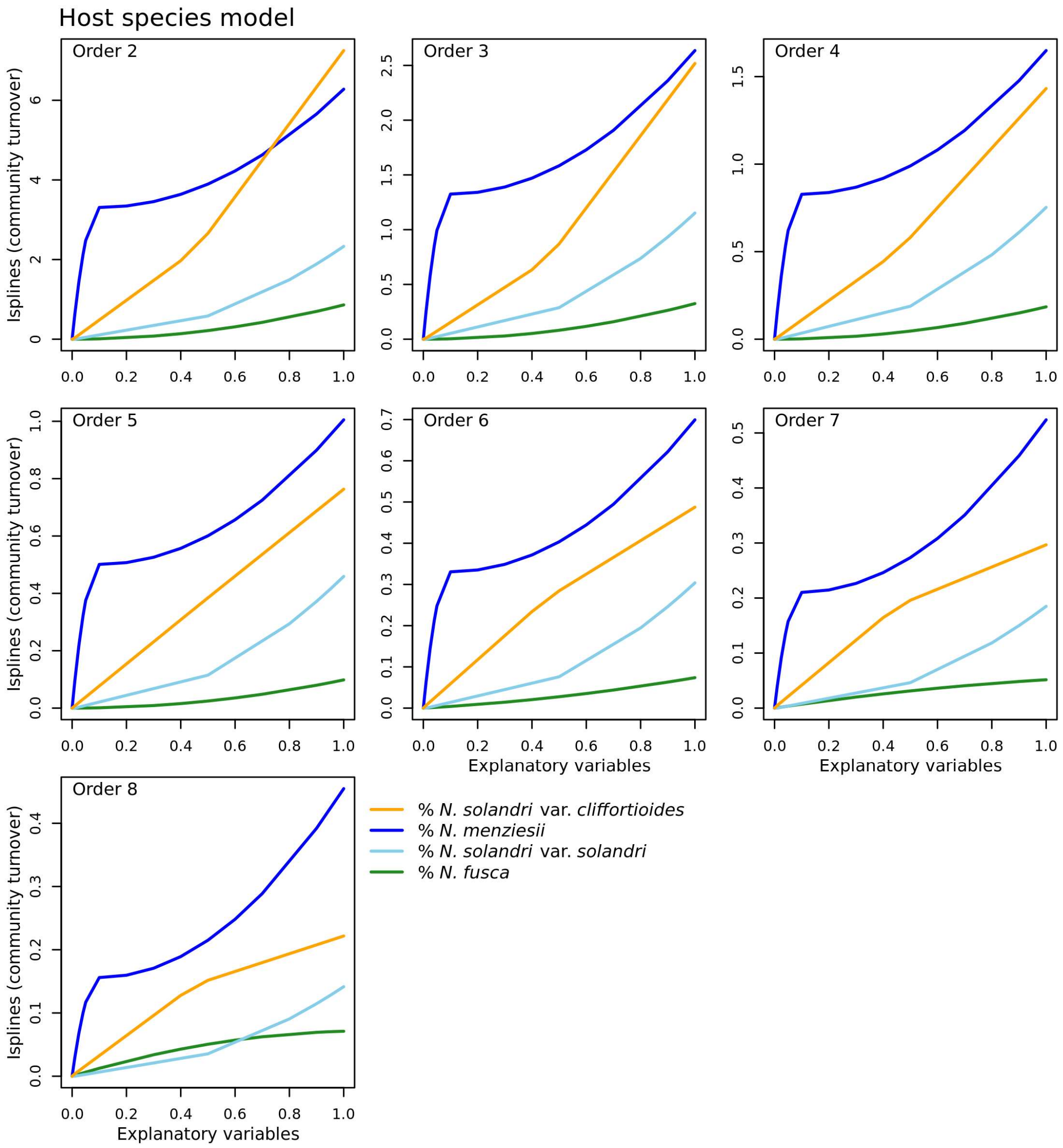


**Figure S5:** Isplines (average of 30 replicates) for the MS-GDM model testing the effect of host variables on fungal community turnover for zeta orders 2 to 8. Variables are rescaled between 0 and 1.

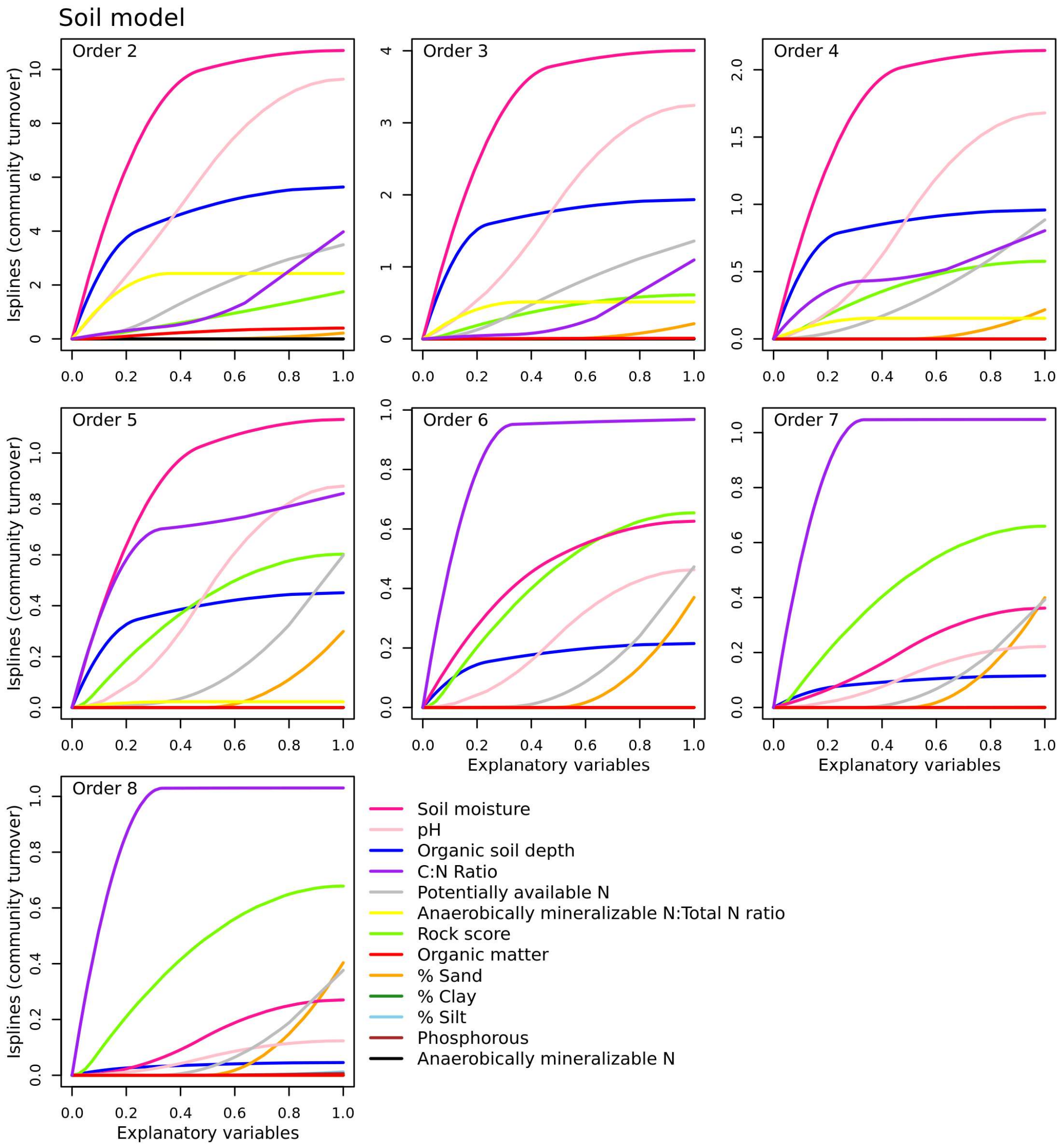


**Figure S6:** Isplines (average of 30 replicates) for the MS-GDM model testing the effect of soil variables on fungal community turnover for zeta orders 2 to 8. Variables are rescaled between 0 and 1.

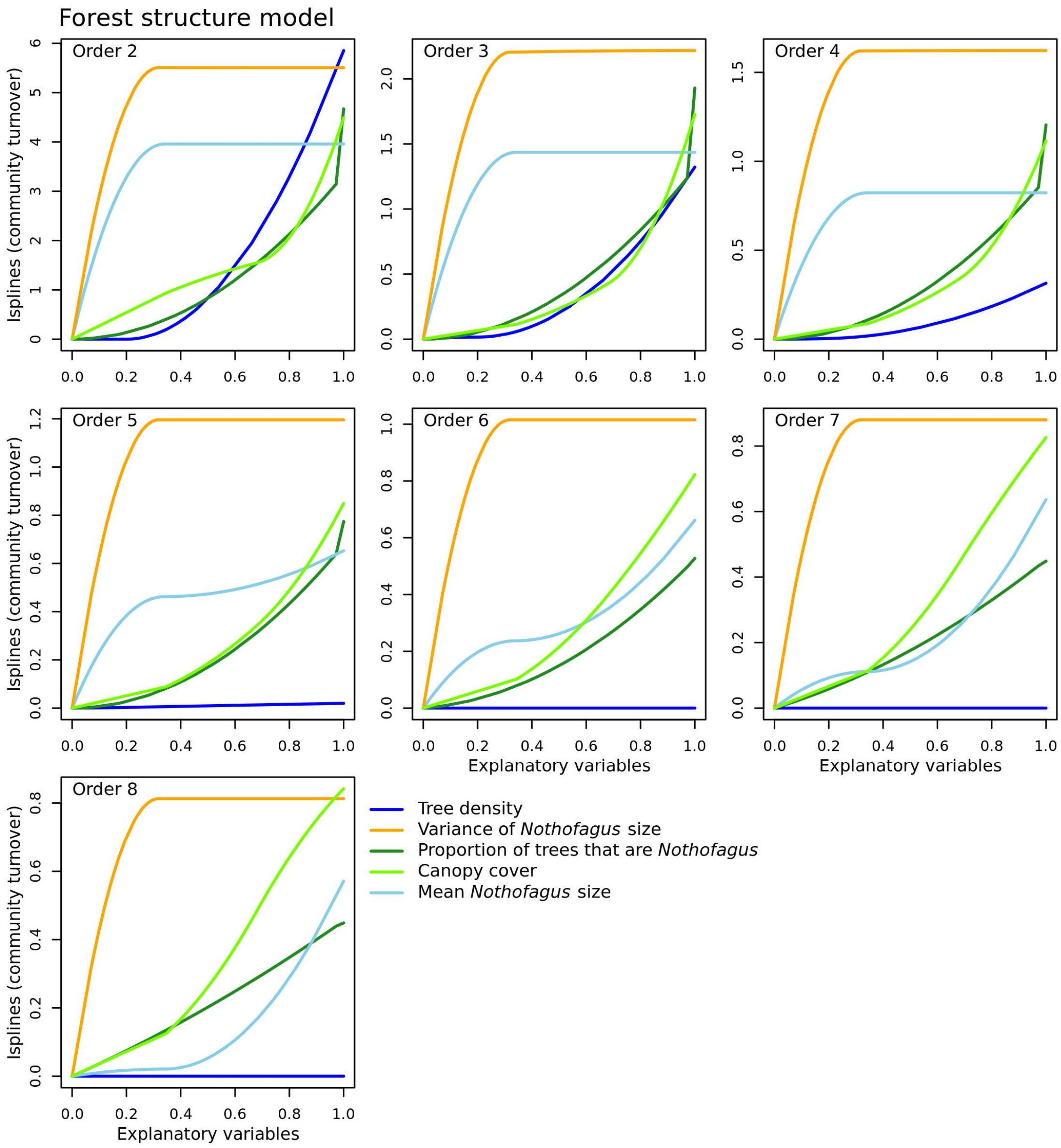


**Figure S7:** Isplines (average of 30 replicates) for the MS-GDM model testing the effect of forest structure variables on fungal community turnover for zeta orders 2 to 8. Variables are rescaled between 0 and 1.

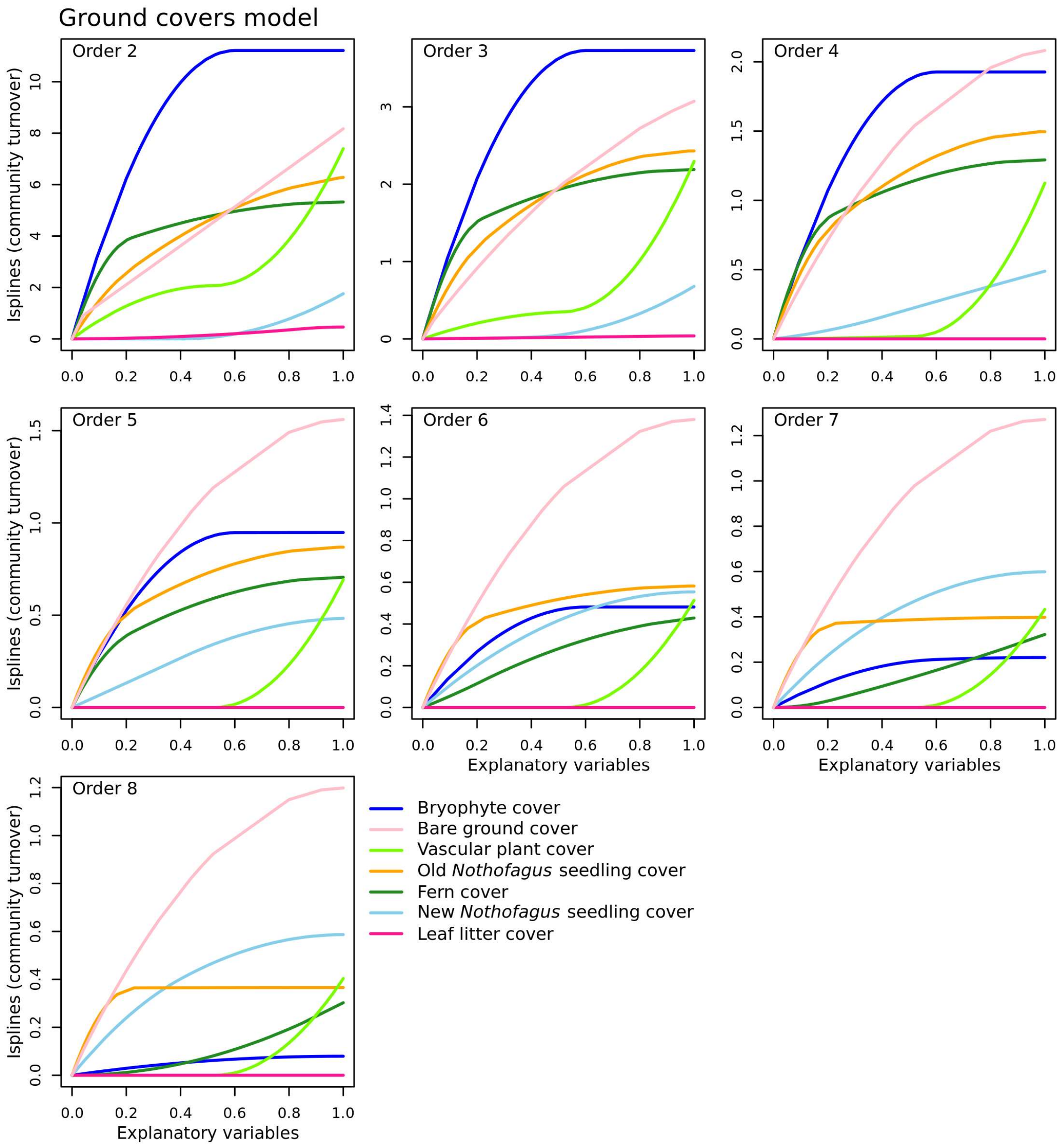


**Figure S8:** Isplines (average of 30 replicates) for the MS-GDM model testing the effect of ground cover variables on fungal community turnover for zeta orders 2 to 8. Variables are rescaled between 0 and 1.

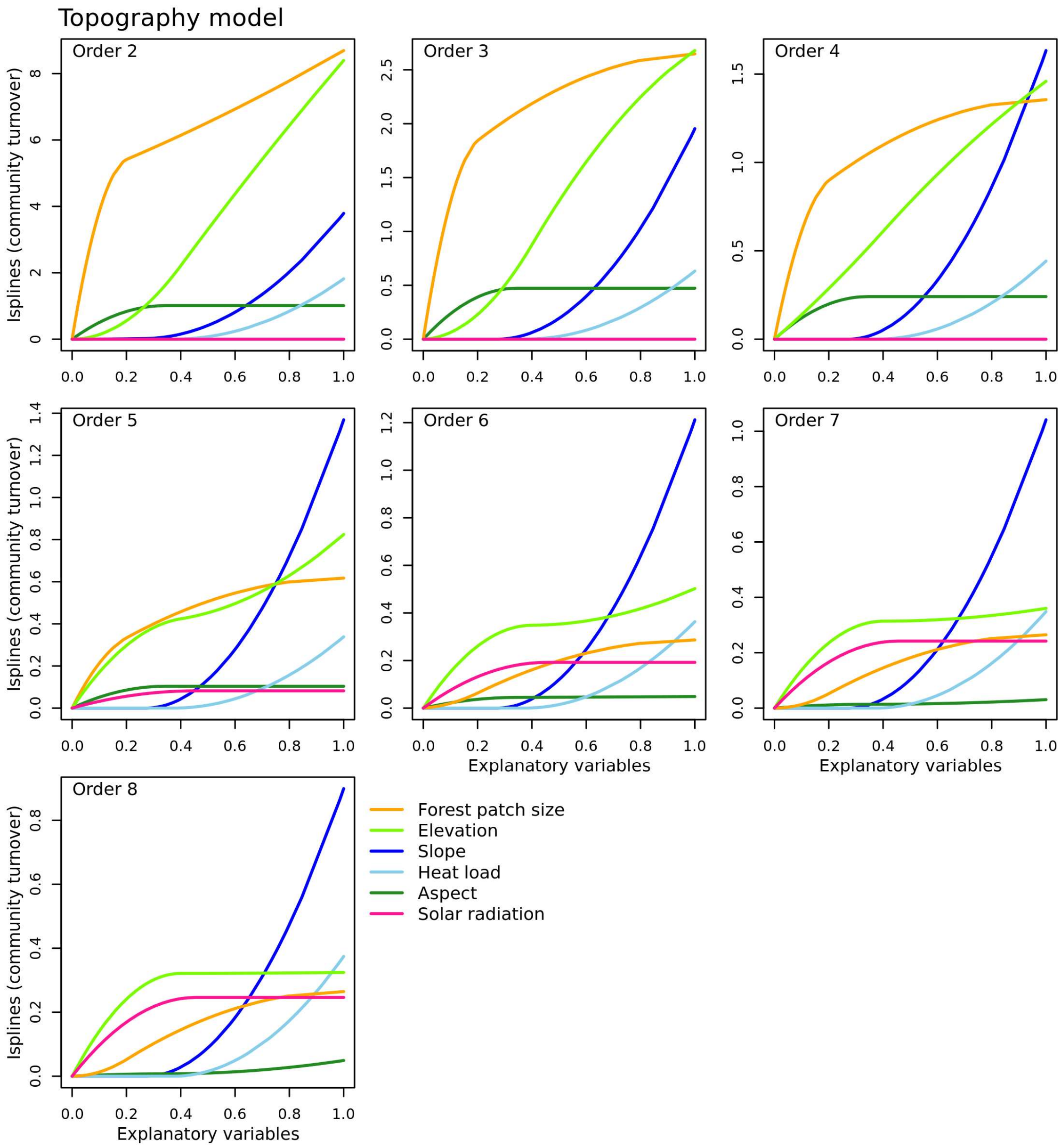


**Figure S9:** Isplines (average of 30 replicates) for the MS-GDM model testing the effect of topographic variables on fungal community turnover for zeta orders 2 to 8. Variables are rescaled between 0 and 1.

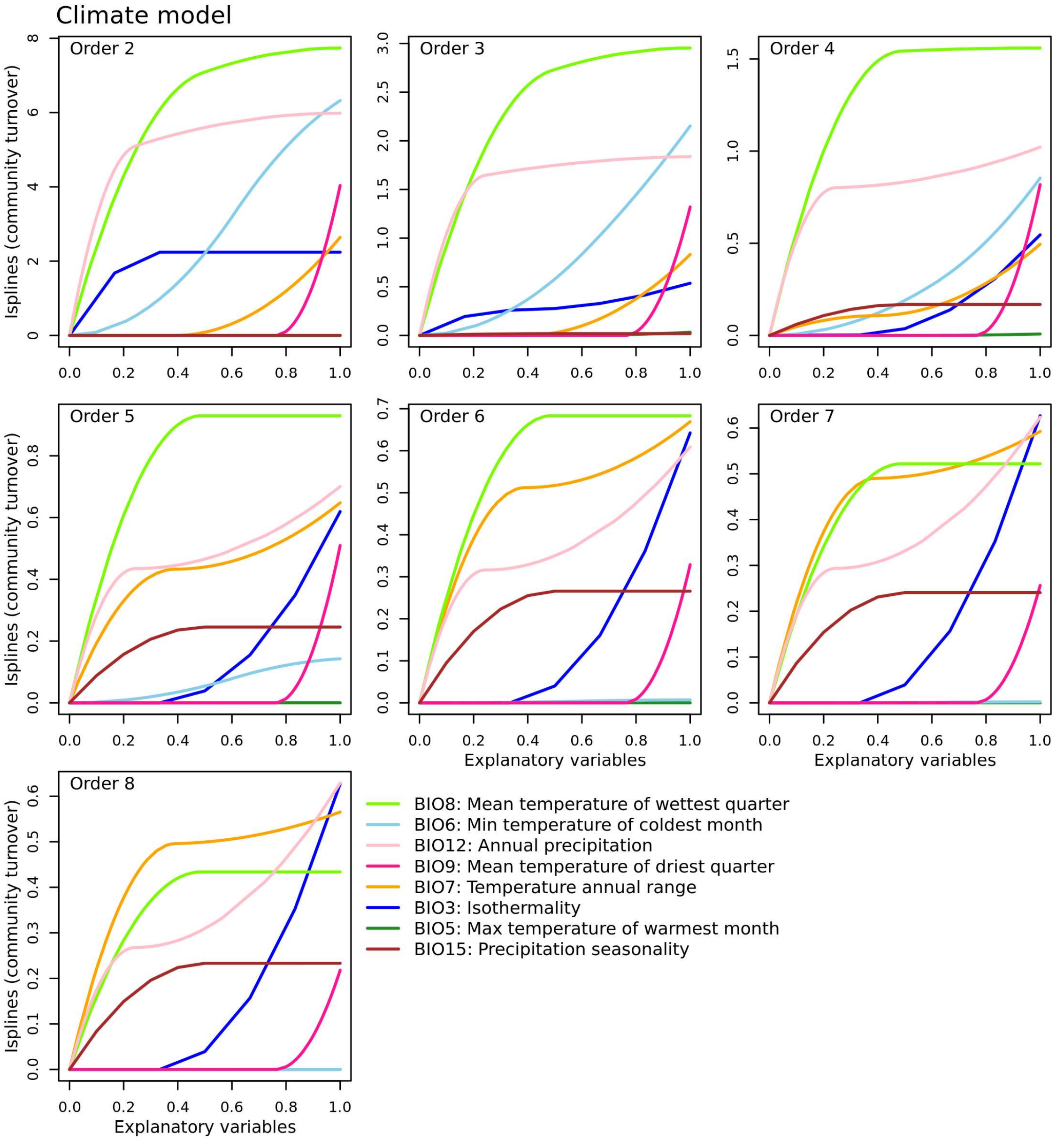


**Figure S10:** Isplines (average of 30 replicates) for the MS-GDM model testing the effect of climatic variables on fungal community turnover for zeta orders 2 to 8. Variables are rescaled between 0 and 1.

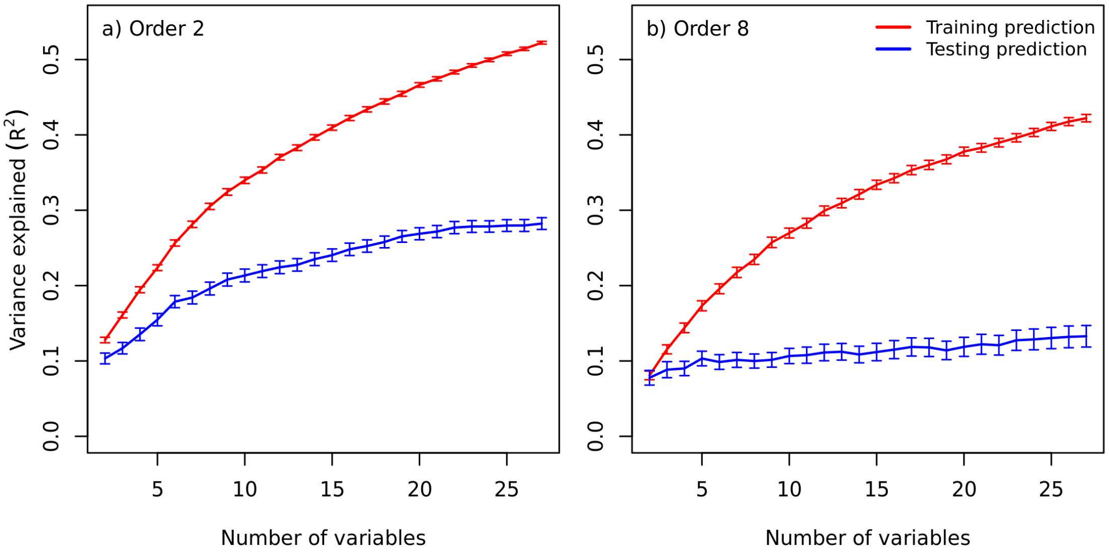


**Figure S11:** The change in variance explained by a) the zeta order 2 and b) the zeta order 8 MS-GDM models with increasing number of predictor variables. Models were built using the training data (see manuscript Section 3.6.2), and then used to predict both the training data values (red) and test data values (blue). Overfitting is evident if the variance explained when predicting the test data (blue) decreases as more variables are included, whilst at the same time the variance explained increases when predicting the training data (red). Therefore, no overfitting was evident, as variance explained for the testing predictions had not begun to decline. Lines show the mean and standard error of 96 replicates of random training and testing divisions for a), and 115 replicates for b). More replicates were used for b) to reduce the standard error and stabilise the pattern.

**Table S2:** The number of ASVs belonging to each genus, the exploration type (from Tedersoo and Smith 2013), and the exploration category used in our analyses. Only genera in bold were included in the analysis (i.e., not those with mixed types). C = contact, SD = short distance, MDF = medium distance fringe, MDM: medium distance mat, MDS medium distance smooth, LD: long distance (see Agerer 2001).

| Genus | Number of ASVs | Exploration type | Category |
|---|---|---|---|
| ***Amanita*** | **34** | **C/SD/MDS** | **Smooth/short** |
| ***Austroboletus*** | **2** | **LD** | **Dense/long** |
| ***Austrogautieria*** | **3** | **MDM** | **Dense/long** |
| ***Austropaxillus*** | **7** | **LD** | **Dense/long** |
| ***Calostoma*** | **10** | **LD** | **Dense/long** |
| *Cenococcum* | 27 | - | - |
| *Chaetospermum* | 15 | - | - |
| *Chlorociboria* | 4 | - | - |
| ***Clavulina*** | **175** | **C** | **Smooth/short** |
| ***Coltricia*** | **4** | **SD** | **Smooth/short** |
| ***Coltriciella*** | **9** | **SD** | **Smooth/short** |
| ***Cortinarius*** | **1403** | **MDF/rarely SD** | **Dense/long** |
| *Cyphellostereum* | 6 | - | - |
| ***Cystangium*** | **48** | **C** | **Smooth/short** |
| ***Dermocybe*** | **11** | **MDF** | **Dense/long** |
| ***Descolea*** | **38** | **SD** | **Smooth/short** |
| ***Descomyces*** | **14** | **SD** | **Smooth/short** |
| ***Dingleya*** | **1** | **C** | **Smooth/short** |
| ***Elaphomyces*** | **4** | **SD** | **Smooth/short** |
| ***Endogone*** | **7** | **SD** | **Smooth/short** |
| ***Fistulinella*** | **4** | **LD** | **Dense/long** |
| ***Gallacea*** | **7** | **MDM** | **Dense/long** |
| *Hebeloma* | 7 | SD/MDF | Mixed |
| ***Hydnellum*** | **4** | **MDM** | **Dense/long** |
| ***Hydnum*** | **9** | **MDS** | **Smooth/short** |
| ***Hysterangium*** | **5** | **MDM** | **Dense/long** |
| ***Inocybe*** | **572** | **SD** | **Smooth/short** |
| ***Laccaria*** | **245** | **SD** | **Smooth/short** |
| *Lachnella* | 1 | - | - |
| ***Lactarius*** | **37** | **C/MDS** | **Smooth/short** |
| *Lactifluus* | 13 | - | - |
| *Lichenomphalia* | 4 | - | - |
| ***Meliniomyces*** | **3** | **SD** | **Smooth/short** |
| *Mycenella* | 13 | - | - |
| *Mycosymbioces* | 5 | - | - |
| *Odontia* | 15 | - | - |
| *Ombrophila* | 1 | - | - |
| *Phaeoclavulina* | 1 | - | - |
| *Phellinotus* | 1 | - | - |
| ***Phellodon*** | **16** | **MDM** | **Dense/long** |
| ***Phylloporus*** | **1** | **LD** | **Dense/long** |
| *Piloderma* | 16 | SD/MDF | Mixed |
| *Pleurella* | 1 | - | - |
| ***Porphyrellus*** | **2** | **LD** | **Dense/long** |
| *Pseudolasiobolus* | 1 | - | - |
| *Ramaria* | 8 | - | - |
| *Rhizocybe* | 2 | - | - |
| ***Rhizopogon*** | **10** | **LD** | **Dense/long** |
| ***Ruhlandiella*** | **819** | **SD** | **Smooth/short** |
| ***Russula*** | **187** | **C** | **Smooth/short** |
| ***Sarcodon*** | **1** | **MDM** | **Dense/long** |
| ***Sebacina*** | **187** | **SD** | **Smooth/short** |
| *Singerocybe* | 1 | - | - |
| ***Suillus*** | **4** | **LD** | **Dense/long** |
| *Tarzetta* | 39 | - | - |
| ***Terfezia*** | **1** | **SD** | **Smooth/short** |
| ***Thelephora*** | **2** | **SD/MDS** | **Smooth/short** |
| ***Tomentella*** | **383** | **C/SD/MDS** | **Smooth/short** |
| ***Tomentellopsis*** | **67** | **MDS** | **Smooth/short** |
| ***Tricholoma*** | **41** | **MDF** | **Dense/long** |

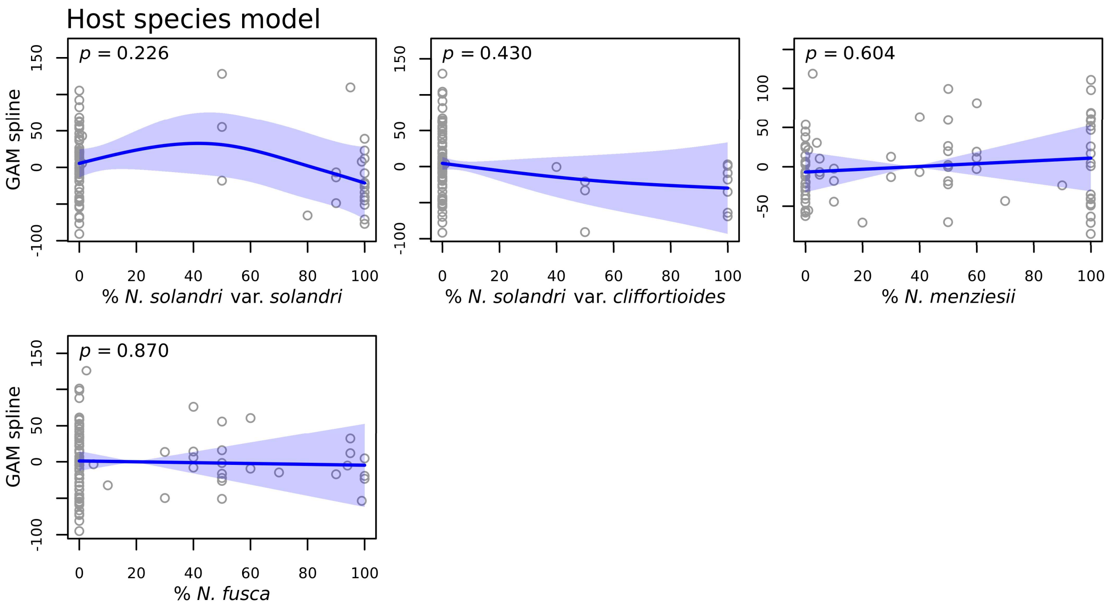


**Figure S12:** The contribution of each variable to the model testing the relationship between host variables and ASV richness. *Nothofagus truncata* was not included due to low replication. Deviance explained = 18.8%, adjusted $R^2$ = 0.13. The blue line shows the fitted smooth curve and shading shows standard error.

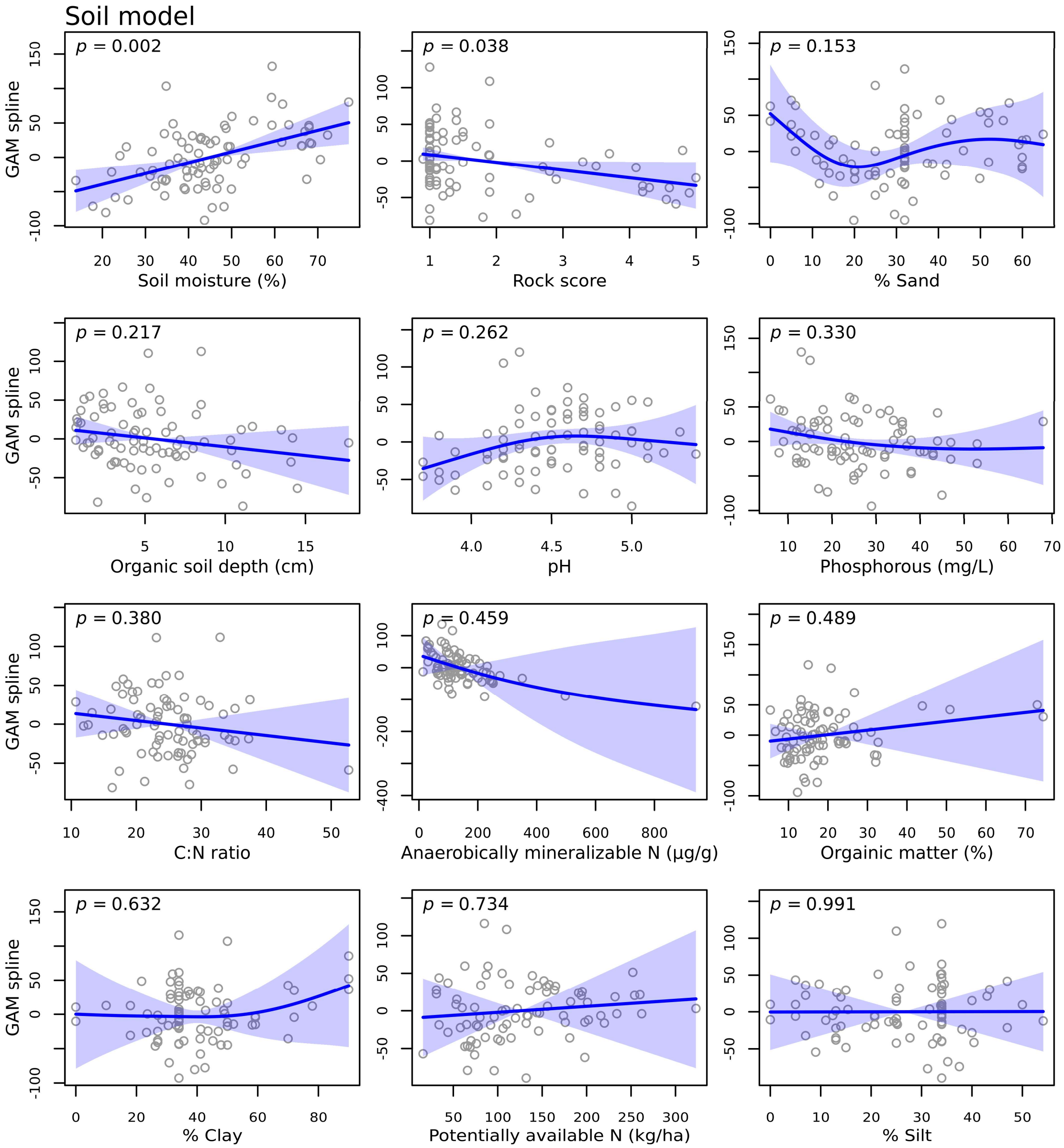


**Figure S13:** The contribution of each variable to the model testing the relationship between soil variables and ASV richness. Deviance explained = 42.4%, adjusted $R^2 = 0.27$. The blue line shows the fitted smooth curve and shading shows standard error. Variables with $p < 0.1$ were used in the combined model.

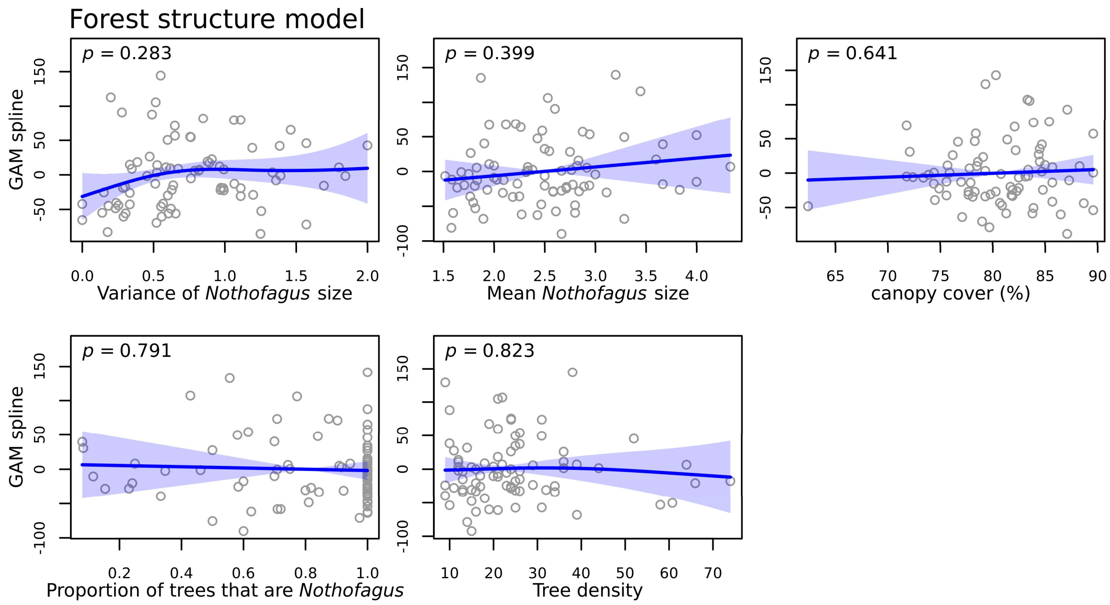


**Figure S14:** The contribution of each variable to the model testing the relationship between forest structure variables and ASV richness. See manuscript Table 1 for an explanation of *Nothofagus* size variable units. Deviance explained = 17.9%, adjusted $R^2 = 0.04$. The blue line shows the fitted smooth curve and shading shows standard error.

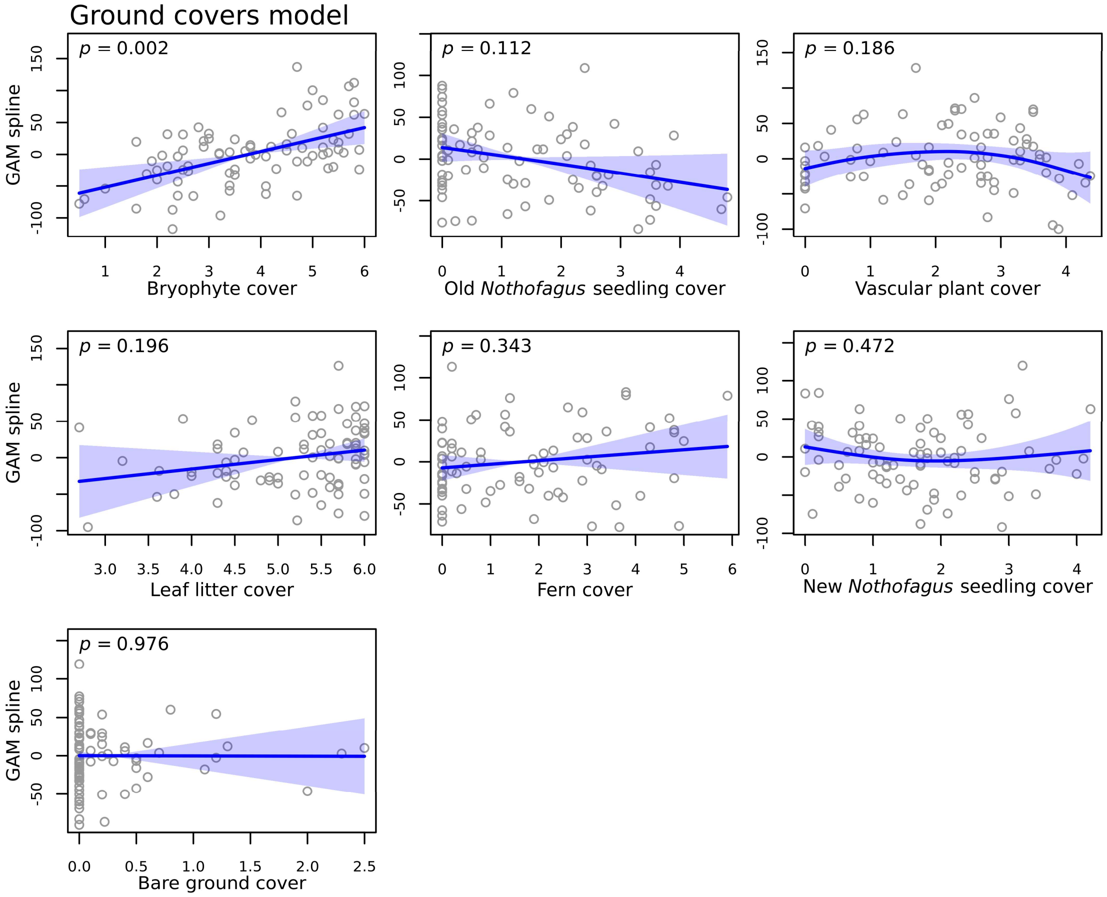


**Figure S15:** The contribution of each variable to the model testing the relationship between ground cover variables and ASV richness. Deviance explained = 31.7%, adjusted $R^2 = 0.23$. The blue line shows the fitted smooth curve and shading shows standard error. Variables with $p < 0.1$ were used in the combined model. Cover variable units are on a Braun-Blanquet scale (see manuscript Section 3.5).

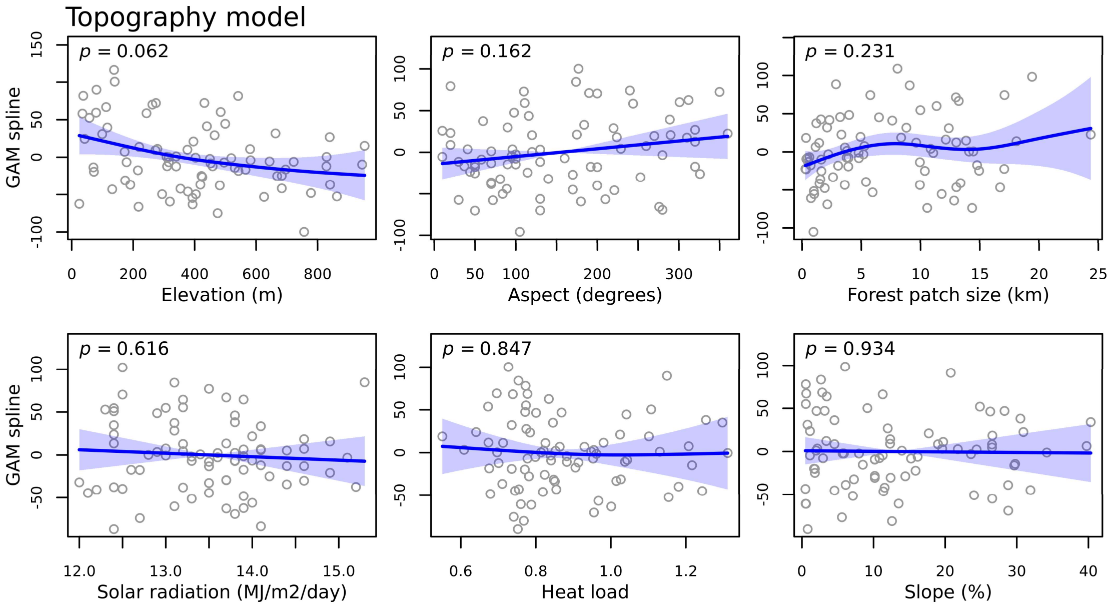


**Figure S16:** The contribution of each variable to the model testing the relationship between topography variables and ASV richness. Deviance explained = 28.8%, adjusted $R^2 = 0.20$. The blue line shows the fitted smooth curve and shading shows standard error. Variables with $p < 0.1$ were used in the combined model.

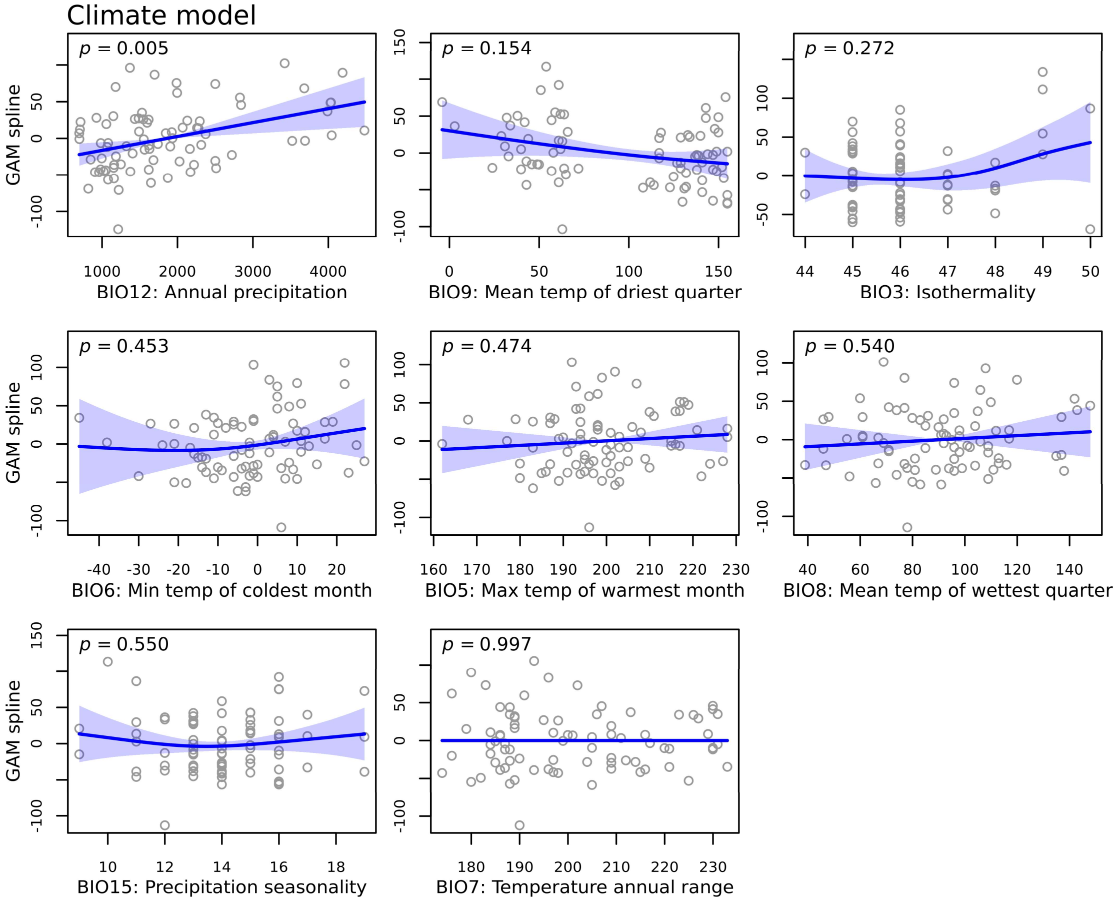


**Figure S17:** The contribution of each variable to the model testing the relationship between climate variables and ASV richness. Units are mm for precipitation variables (BIO12, BIO15), and °C multiplied by 10 for temperature variables (BIO5, BIO6, BIO7, BIO8, BIO9). BIO3 (isothermality) is the ratio of mean diurnal temperature range to annual temperature range. Deviance explained = 28.8%, adjusted $R^2$ = 0.20. The blue line shows the fitted smooth curve and shading shows standard error. Variables with $p < 0.1$ were used in the combined model.

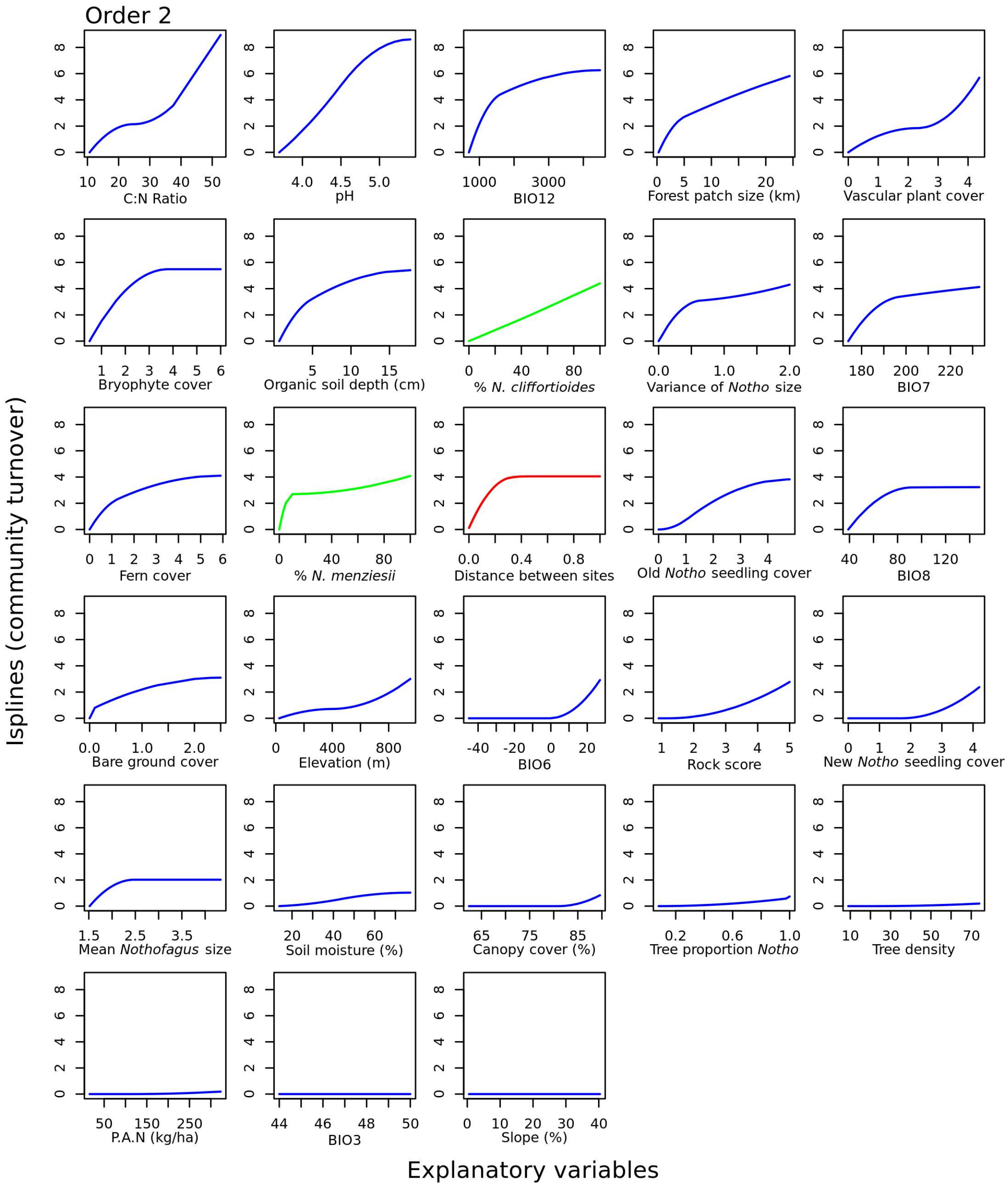


**Figure S18:** Isplines for all 27 variables included in the combined zeta order 2 MS-GDM model. The amplitude of each ispline indicates the relative contribution of each variable to fungal community turnover, allowing the relative importance of distance between sites (red) and *Nothofagus* host species (green) to be compared to other environmental variables (blue). The slope of the ispline indicates the rate of community turnover occurring at different points along the environmental gradient. P.A.N = Potentially available nitrogen, Tree proportion *Notho* = Proportion of trees that are *Nothofagus*, BIO3 = Isothermality, BIO6 = Minimum temperature of coldest month (°C × 10), BIO7 = Temperature annual range (°C × 10), BIO 8 = Mean temperature of wettest quarter (°C × 10), BIO12 = Annual precipitation (mm). Distance and *Nothofagus* host species effects are scaled between 0 and 1, see manuscript Table 2 for an explanation of cover and *Nothofagus* size variable units.

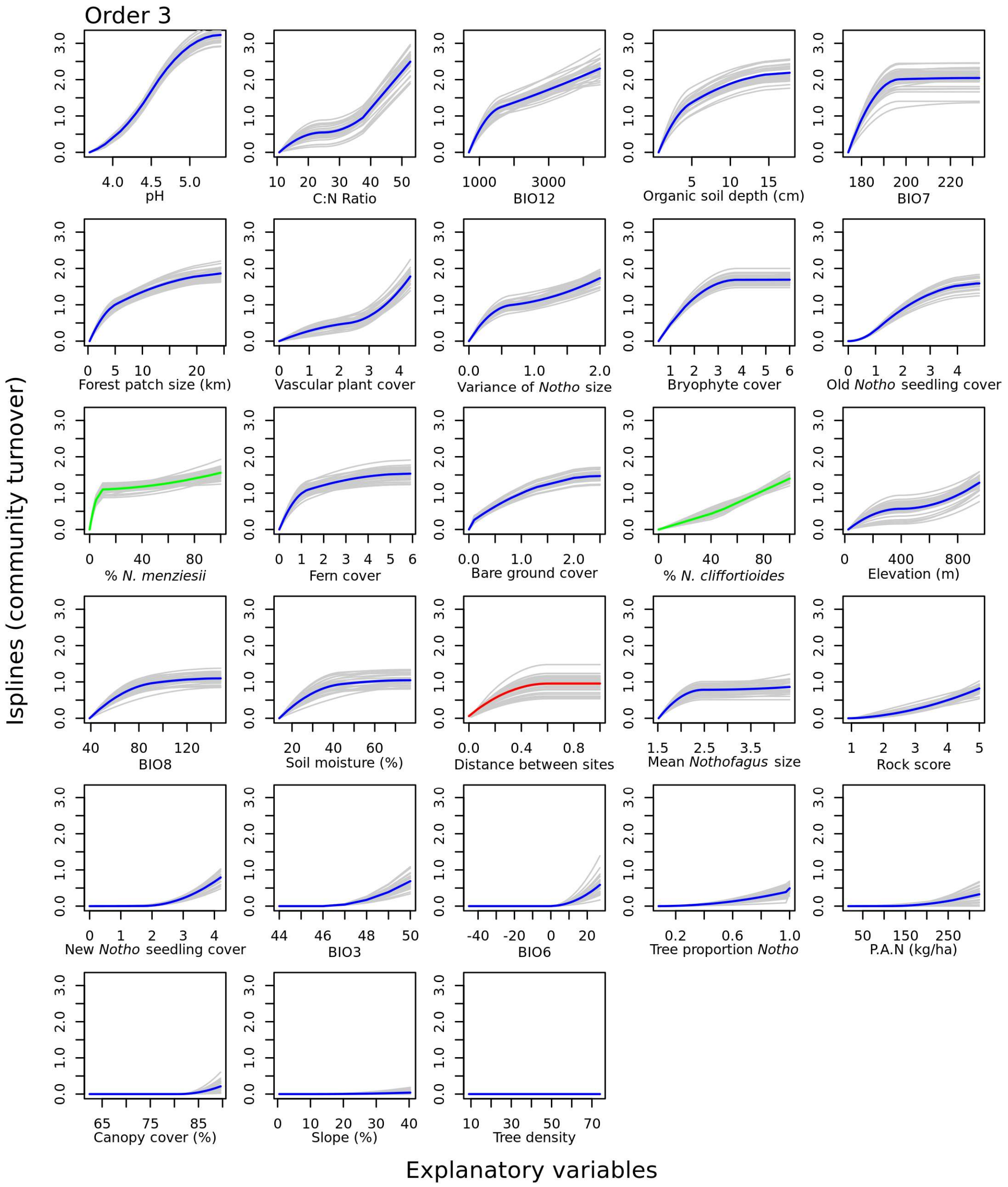


**Figure S19:** Isplines for all 27 variables included in the combined zeta order 3 MS-GDM model. The amplitude of each ispline indicates the relative contribution of each variable to fungal community turnover, allowing the relative importance of distance between sites (red) and *Nothofagus* host species (green) to be compared to other environmental variables (blue). The slope of the ispline indicates the rate of community turnover occurring at different points along the environmental gradient. Coloured lines are the average of 30 replicates (shown in grey). P.A.N = Potentially available nitrogen, Tree proportion *Notho* = Proportion of trees that are *Nothofagus*, BIO3 = Isothermality, BIO6 = Minimum temperature of coldest month (°C × 10), BIO7 = Temperature annual range (°C × 10), BIO 8 = Mean temperature of wettest quarter (°C × 10), BIO12 = Annual precipitation (mm). Distance and *Nothofagus* host species effects are scaled between 0 and 1, see manuscript Table 2 for an explanation of cover and *Nothofagus* size variable units.

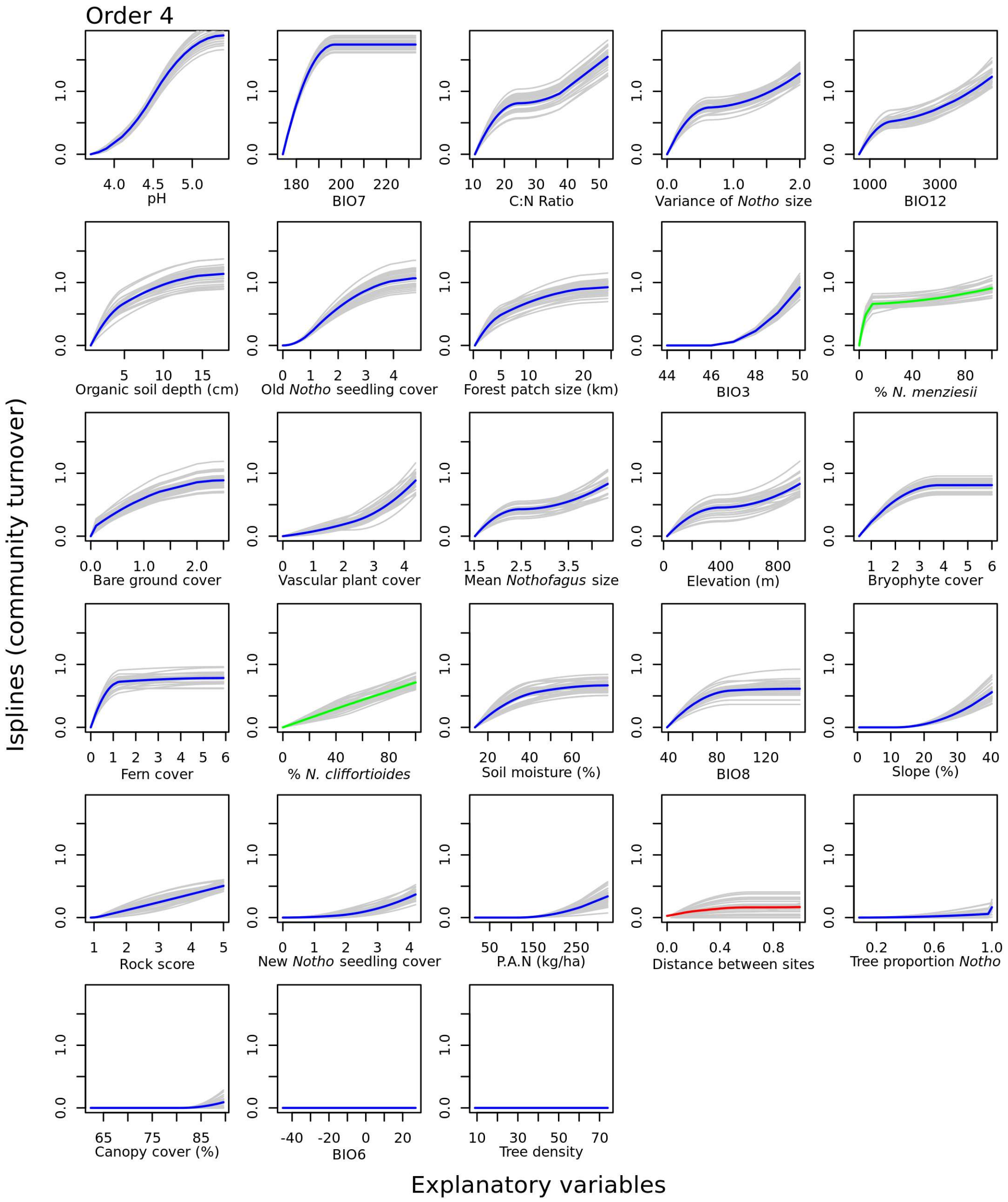


**Figure S20:** Isplines for all 27 variables included in the combined zeta order 4 MS-GDM model. The amplitude of each ispline indicates the relative contribution of each variable to fungal community turnover, allowing the relative importance of distance between sites (red) and *Nothofagus* host species (green) to be compared to other environmental variables (blue). The slope of the ispline indicates the rate of community turnover occurring at different points along the environmental gradient. Coloured lines are the average of 30 replicates (shown in grey). P.A.N = Potentially available nitrogen, Tree proportion *Notho* = Proportion of trees that are *Nothofagus*, BIO3 = Isothermality, BIO6 = Minimum temperature of coldest month (°C × 10), BIO7 = Temperature annual range (°C × 10), BIO 8 = Mean temperature of wettest quarter (°C × 10), BIO12 = Annual precipitation (mm). Distance and *Nothofagus* host species effects are scaled between 0 and 1, see manuscript Table 2 for an explanation of cover and *Nothofagus* size variable units.

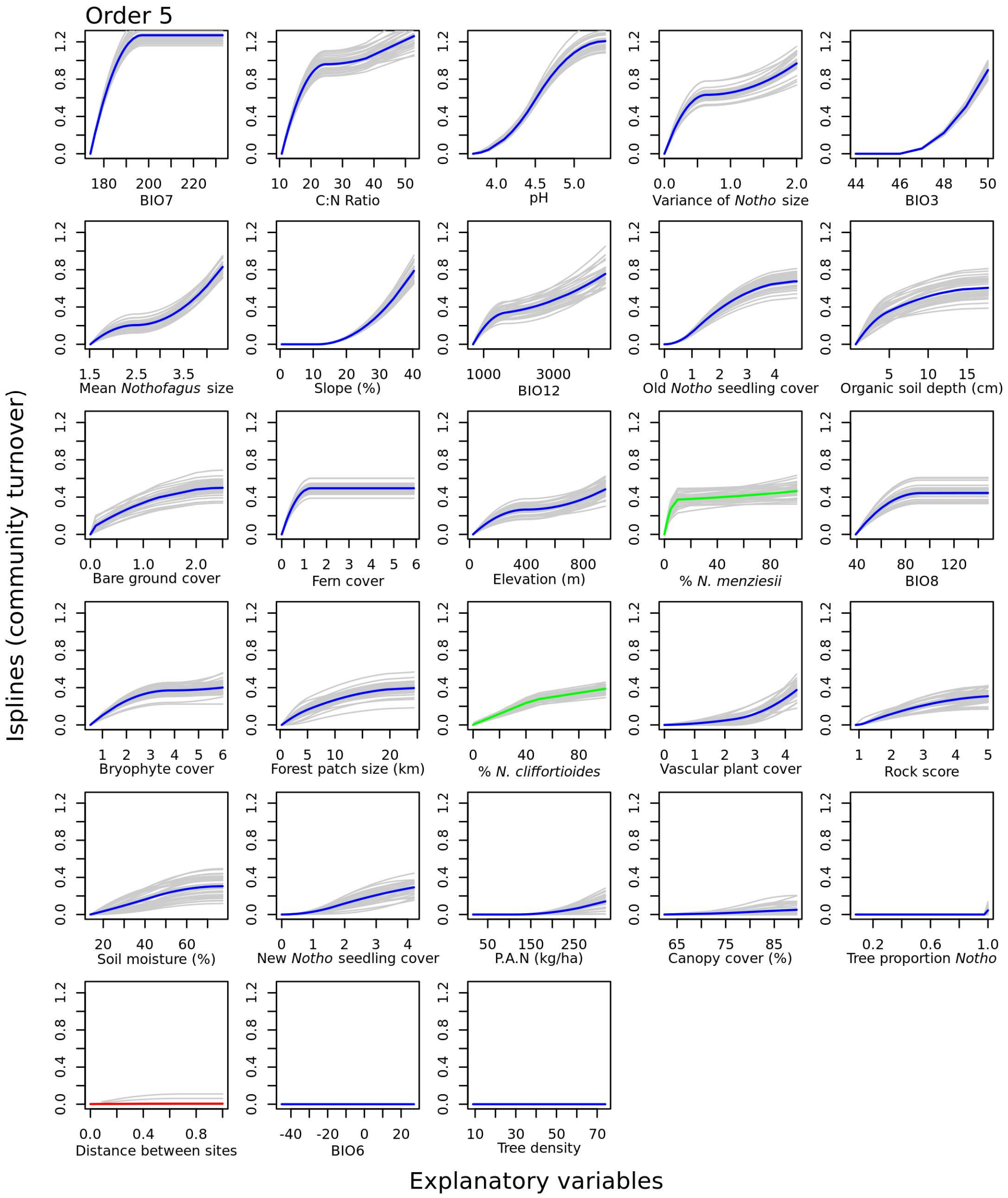


**Figure S21:** Isplines for all 27 variables included in the combined zeta order 5 MS-GDM model. The amplitude of each ispline indicates the relative contribution of each variable to fungal community turnover, allowing the relative importance of distance between sites (red) and *Nothofagus* host species (green) to be compared to other environmental variables (blue). The slope of the ispline indicates the rate of community turnover occurring at different points along the environmental gradient. Coloured lines are the average of 30 replicates (shown in grey). P.A.N = Potentially available nitrogen, Tree proportion *Notho* = Proportion of trees that are *Nothofagus*, BIO3 = Isothermality, BIO6 = Minimum temperature of coldest month (°C × 10), BIO7 = Temperature annual range (°C × 10), BIO 8 = Mean temperature of wettest quarter (°C × 10), BIO12 = Annual precipitation (mm). Distance and *Nothofagus* host species effects are scaled between 0 and 1, see manuscript Table 2 for an explanation of cover and *Nothofagus* size variable units.

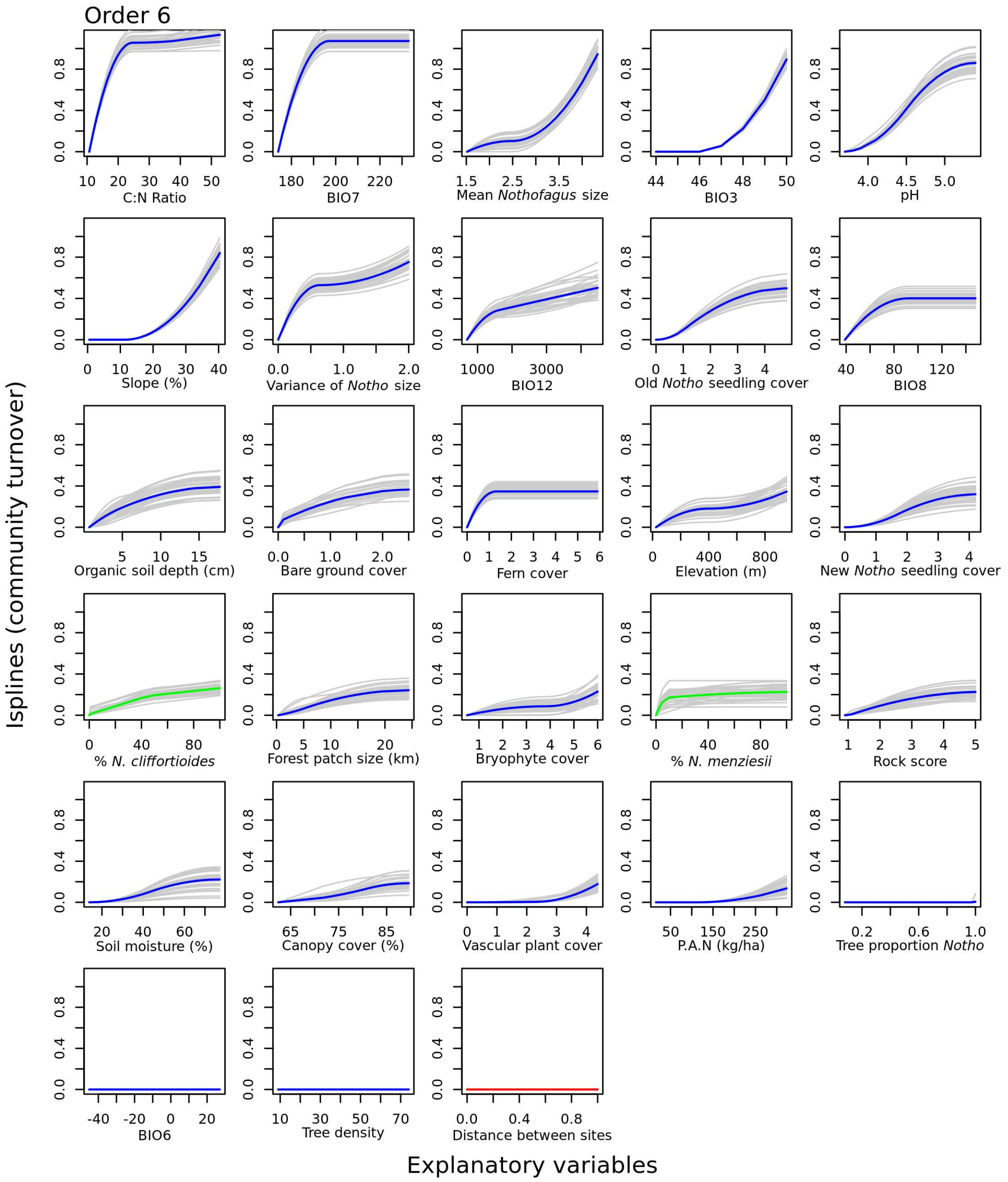


**Figure S22:** Isplines for all 27 variables included in the combined zeta order 6 MS-GDM model. The amplitude of each ispline indicates the relative contribution of each variable to fungal community turnover, allowing the relative importance of distance between sites (red) and *Nothofagus* host species (green) to be compared to other environmental variables (blue). The slope of the ispline indicates the rate of community turnover occurring at different points along the environmental gradient. Coloured lines are the average of 30 replicates (shown in grey). P.A.N = Potentially available nitrogen, Tree proportion *Notho* = Proportion of trees that are *Nothofagus*, BIO3 = Isothermality, BIO6 = Minimum temperature of coldest month (°C × 10), BIO7 = Temperature annual range (°C × 10), BIO 8 = Mean temperature of wettest quarter (°C × 10), BIO12 = Annual precipitation (mm). Distance and *Nothofagus* host species effects are scaled between 0 and 1, see manuscript Table 2 for an explanation of cover and *Nothofagus* size variable units.

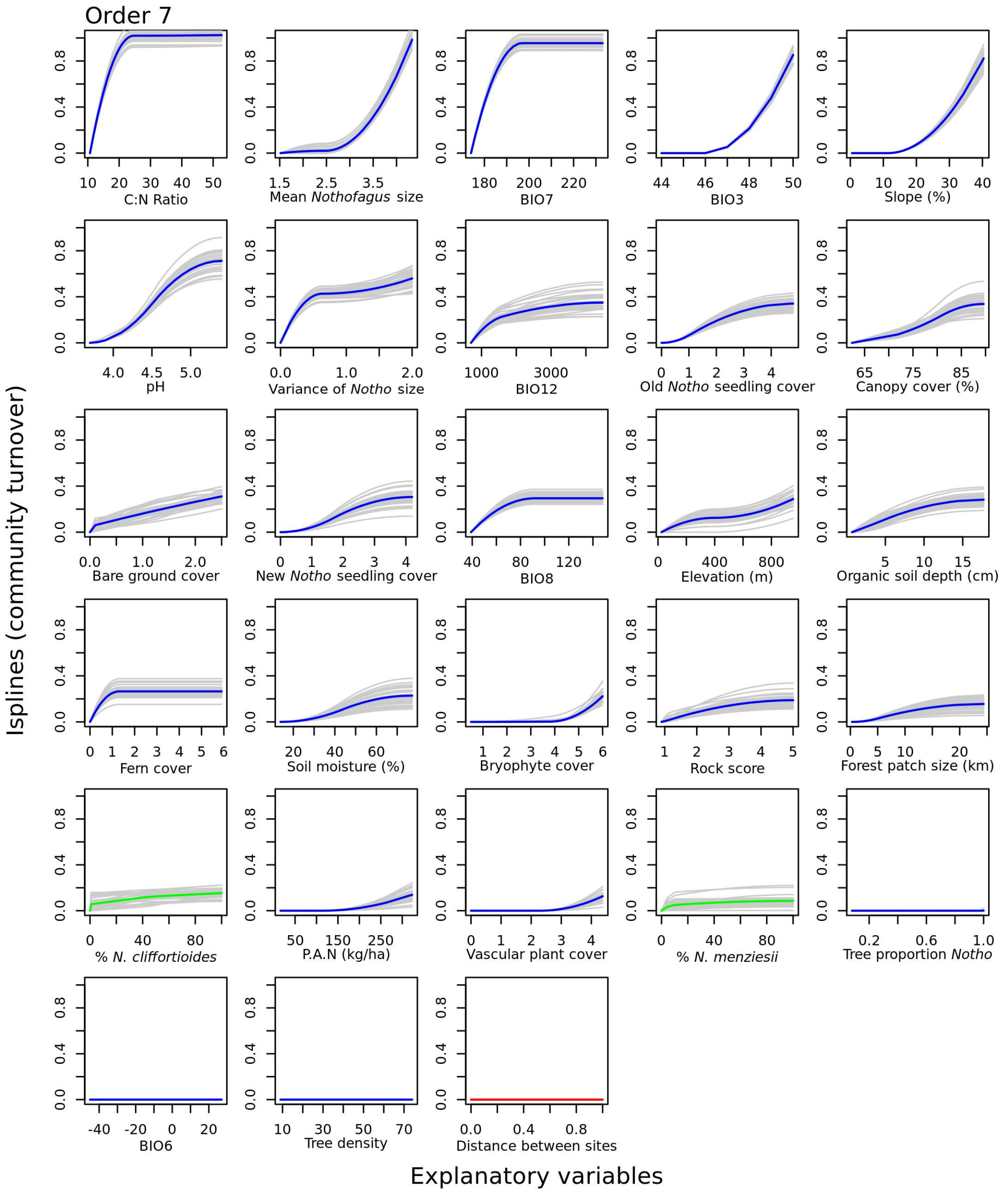


**Figure S23:** Isplines for all 27 variables included in the combined zeta order 7 MS-GDM model. The amplitude of each ispline indicates the relative contribution of each variable to fungal community turnover, allowing the relative importance of distance between sites (red) and *Nothofagus* host species (green) to be compared to other environmental variables (blue). The slope of the ispline indicates the rate of community turnover occurring at different points along the environmental gradient. Coloured lines are the average of 30 replicates (shown in grey). P.A.N = Potentially available nitrogen, Tree proportion *Notho* = Proportion of trees that are *Nothofagus*, BIO3 = Isothermality, BIO6 = Minimum temperature of coldest month (°C × 10), BIO7 = Temperature annual range (°C × 10), BIO 8 = Mean temperature of wettest quarter (°C × 10), BIO12 = Annual precipitation (mm). Distance and *Nothofagus* host species effects are scaled between 0 and 1, see manuscript Table 2 for an explanation of cover and *Nothofagus* size variable units.

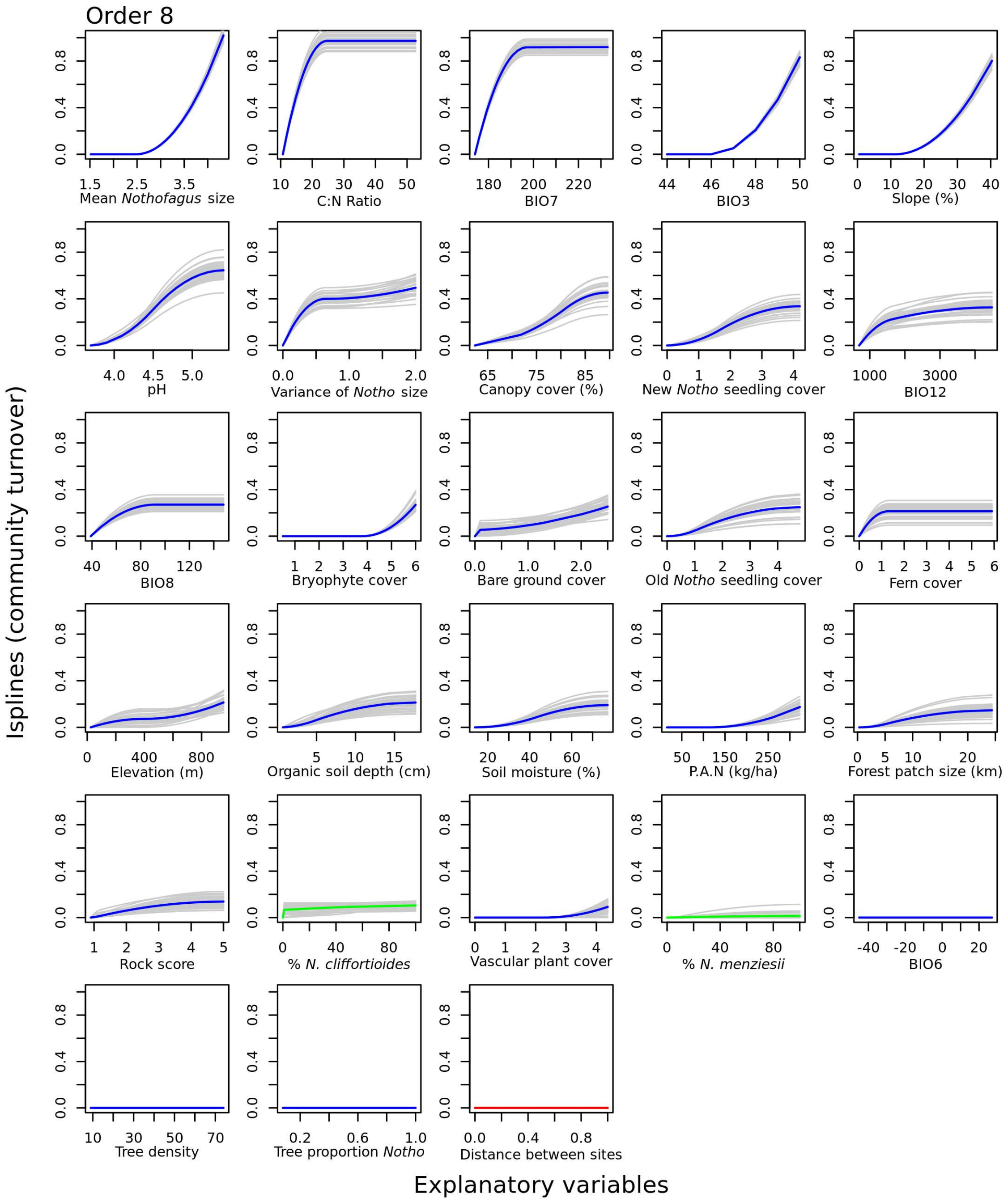


**Figure S24:** Isplines for all 27 variables included in the combined zeta order 8 MS-GDM model. The amplitude of each ispline indicates the relative contribution of each variable to fungal community turnover, allowing the relative importance of distance between sites (red) and *Nothofagus* host species (green) to be compared to other environmental variables (blue). The slope of the ispline indicates the rate of community turnover occurring at different points along the environmental gradient. Coloured lines are the average of 30 replicates (shown in grey). P.A.N = Potentially available nitrogen, Tree proportion *Notho* = Proportion of trees that are *Nothofagus*, BIO3 = Isothermality, BIO6 = Minimum temperature of coldest month (°C × 10), BIO7 = Temperature annual range (°C × 10), BIO 8 = Mean temperature of wettest quarter (°C × 10), BIO12 = Annual precipitation (mm). Distance and *Nothofagus* host species effects are scaled between 0 and 1, see manuscript Table 2 for an explanation of cover and *Nothofagus* size variable units.
